# Damsels in a hidden colour: development of ultraviolet sensitivity and colour patterns in damselfishes (Pomacentridae)

**DOI:** 10.1101/2024.08.22.609266

**Authors:** Valerio Tettamanti, N. Justin Marshall, Karen L. Cheney, Fabio Cortesi

## Abstract

Damselfishes (Pomacentridae) are widespread and highly abundant on tropical coral reefs. They exhibit diverse body colouration within and between the ∼250 species and across ontogenetic stages. In addition to human visible colours (i.e., 400-700 nm), most adult damselfishes reflect ultraviolet (UV, 300-400 nm) colour patches. UV sensitivity and UV colour signals are essential for feeding and form the basis for a secret communication channel invisible to the many UV-blind predatory fish on the reef; however, how these traits develop across ontogenetic stages, and their distribution across the damselfish family is poorly characterised. Here, we used UV photography, phylogenetic reconstructions of opsin genes, differential gene expression analysis (DGE) of retinal samples, to investigate the development of UV vision and colour patterns in three ontogenetic stages (pre-settlement larval, juvenile, and adult) of eleven damselfish species. Using DGE, we found similar gene expression between juveniles and adults, which strongly differed from larvae. All species and all stages expressed at least one UV-sensitive *sws1* opsin gene. However, UV body colour patterns only started to appear at the juvenile stage. Moreover, *Pomacentrus* species displayed highly complex UV body patterns that were correlated with the expression of two *sws1* copies. This could mean that some damselfishes can discriminate colours that change only in their UV component. We demonstrate dramatic shifts in both UV sensitivity and UV colouration across the development stages of damselfish, while highlighting the importance of considering ontogeny when studying the coevolution of visual systems and colour signals.

## Introduction

Animals use colour vision for critical tasks such as feeding, mating, predator avoidance and navigation (Cronin et al., 2014). To perform these tasks efficiently in highly variable light environments, such as those on tropical coral reefs, fishes have evolved visual systems to perceive short-wavelengths of light (ultraviolet UV, ∼ 350 nm) up to longer-wavelengths (red, ∼ 600 nm) (reviewed by Cortesi et al., 2020). Damselfishes (Pomacentridae), one of the most prevalent reef fish families, display a high species diversity, a variety of ecologies, and differ widely in colouration and morphology (Allen, 1991). Damselfish visual systems also vary significantly as they differ in visual gene (opsin) expression and structure (Hofmann et al., 2012; Stieb et al., 2024, 2023, 2019, 2017, 2016). A notable feature of damselfishes is their ability to perceive UV wavelengths, facilitated by a UV-transmitting lens and cornea (Siebeck and Marshall, 2007, 2001), and a UV-sensitive photoreceptor that expresses the short-wavelength-sensitive 1 (*sws1*) opsin gene (Job and Bellwood, 2007; Mitchell et al., 2024, 2023; Powell et al., 2021; Siebeck et al., 2010).

In teleost fishes, five different types of visual opsin genes are found. Rod photoreceptors express the rhodopsin (*rh1*). In contrast, cone photoreceptors express *sws1* and *sws2*, mid-wavelength sensitive rhodopsin-like 2 (*rh2*), and long-wavelength sensitive (*lws*) opsin genes (Carleton et al., 2020; Musilova et al., 2019). Opsin genes in teleosts have experienced a dynamic evolutionary history affected by gene duplications, gene losses, sequence modifications, and gene conversion (Carleton et al., 2020; Cortesi et al., 2015; Mitchell et al., 2021; Musilova et al., 2021, 2019; Musilova and Cortesi, 2023). The latter describes a process by which an unequal crossing over during meiosis results in a unidirectional transfer of genetic information and consequential identical residues in different genes (Chen et al., 2007; Holliday, 1964). Fishes can also change the expression of opsin genes over development or shorter timescales, and some species have been found to co-express multiple opsins in the same photoreceptor (Cortesi et al., 2016; Dalton et al., 2014; Savelli et al., 2018; Torres-Dowdall et al., 2021). Moreover, some fishes can also convert their chromophore from A_1_-derived to A_2_-derived using the CYP27C1 enzyme, switching the sensitivity of visual pigments to longer wavelengths (Enright et al., 2015). These processes allow visual adaptations to different light environments, behaviours, and ecologies across generations or within the lifetime of a species (Carleton et al., 2020).

Damselfishes express different sets of these visual opsin genes, with some species only expressing three and others expressing up to six cone opsin copies (Mitchell et al., 2021; Stieb et al., 2024, 2023, 2019, 2017, 2016). In adult damselfishes, single cones have been found to express the short-wavelength-sensitive opsin genes (*sws1* and *sws2b*), and the double cones (two fused single cones; Walls, 1942) to express the mid- and long-wavelength-sensitive genes (*rh2*s and *lws*). The adults of some, but not all, damselfish species have also been found to tune opsin gene expression over short periods (weeks to months), with depth, and between seasons to adapt to changes in their light environments (Luehrmann et al., 2018; Stieb et al., 2016). Moreover, species-specific ecologies and colouration seem to influence opsin gene expression: longer wavelength sensitivity occurs in herbivorous damselfishes and is more pronounced in species with red colouration, while shorter wavelength sensitivity correlates with UV body colour patterns (Stieb et al., 2024, 2023, 2019, 2017, 2016). Behavioural studies have found that these small fishes use UV signals to communicate with con- and hetero-specifics (Mitchell et al., 2023; Siebeck et al., 2010). Because larger predatory fish are UV-blind, this has led to the hypothesis that damselfishes use UV vision and colouration as a ‘private communication channel’ (Siebeck, 2004; Siebeck et al., 2010; Stieb et al., 2017). Moreover, in freshwater fishes, UV patterns correlate to sexual selection (Macías Garcia and de Perera, 2002; Smith et al., 2002). Some damselfish species, particularly anemonefishes, also express multiple *sws1* copies (Mitchell et al., 2021; Stieb et al., 2024). The *sws1* gene duplication seems to have occurred independently at least twice in the damselfish family. One duplication occurred in the Pomacentrinae subfamily and has been dated to the last common ancestor of the *Pomacentrus*, *Neopomacentrus* and *Amphiprion* genera (termed Pomacentrinae 3, 4 and 5 in McCord et al., 2021 (Stieb et al., 2024). A second, species-specific duplication was discovered in the genome of *Chromis chromis* (Musilova et al., 2019). The two Pomacentrinae copies cluster in short (*sws1*α; λ_max_ 356-362 nm) and longer sensitive (*sws1*β; λ_max_ 368-370 nm) clades depending on changes in key amino acid sites of the opsin protein at positions 114 and 118 (Mitchell et al., 2021; Stieb et al., 2024). However, how widespread the expression of these copies is and their function remained unclear.

While there have been several studies on adult damselfish vision (e.g., Luehrmann et al., 2018; Siebeck et al., 2010, 2010; Stieb et al., 2017), very little is known about the visual systems of earlier developmental stages. Studying ontogenetic changes in coral reef fish vision is of great importance to better understand how they visually adapt to face diverse challenges encountered through their life stages (e.g. predation and competition for food and shelter, see Anderson, 2001; Carr et al., 2002; Hixon and Carr, 1997). During the pelagic larval stage, damselfishes are transparent and feed on zooplankton (Sampey et al., 2007). Most zooplankton either reflect or absorb UV light, and it has been shown that fishes, including salmonids (Flamarique, 2000), zebrafish (Baden, 2021), cichlids (Jordan et al., 2004), and larval damselfish (Job and Shand, 2001), use UV vision to spot them against the UV-lit background in the water column. Once damselfishes settle on the reef and metamorphose into juveniles, they show various feeding ecologies and colour patterns, likely to contain UV colours. Often, damselfish colours and patterns, at least in the human visible, change again when they turn into adults (Frédérich and Parmentier, 2016).

Skin reflectance measurements using spectrophotometers have commonly been used to assess the colour of damselfishes, especially in the UV (Marshall et al., 2019). However, these measurements do not contain information about the spatial distribution of UV patterns. Only in two sister species of damsefish (*Pomacentrus amboinensis* and *P. moluccensis*) (Siebeck et al., 2010) has UV photography been used to assess the nature of UV patterns. In theseb, the UV patterns show a high degree of complexity with differently shaped dotted and lined motifs on the operculum and simpler motifs on the fins and the body that the fish use to distinguish individuals, similar to a human fingerprint (Gagliano et al., 2015; Siebeck, 2014, 2004; Siebeck et al., 2010).

In this study, we hypothesised that complex UV colour patterns with variations in shape and structure are widespread in the damselfish family. However, these patterns emerge only in later developmental stages, reflecting the ontogenetic change in colour communications of these species. Moreover, because several adult damselfishes have been found to express two *sws1* opsins with different absorption maxima, it is possible that these fishes can distinguish between colours that only differ in UV wavelengths. Colour discrimination between UV wavelengths alone has been shown in butterflies (Finkbeiner and Briscoe, 2021) and in mantis shrimp (Thoen et al., 2014), and besides needing multiple photoreceptors with different UV sensitivity, it also necessitates UV colours that have dissimilar peaks in the spectral curve. Hence, we predicted the expression of two *sws1* opsins to correlate with the occurrence of UV patterns that differ in structural complexity and spectral reflectance. To test our hypotheses, we first used standardised photography in the human visible (i.e., RGB) and UV to correlate expression changes with shifts in ecology and colouration, focusing on UV complex patterns, *sws1* expression and considering the damselfish phylogeny. We then used comparative transcriptomics to investigate ontogenetic changes in opsin gene expression in eleven damselfish species from two (Pomacentrinae, Chrominae) of the four damselfish subfamilies (McCord et al., 2021).

## Methods

### Specimen collection

Eleven damselfish (Pomacentridae) species (subfamily Chrominae*: Chromis atripectoralis*, *Dascyllus aruanus*; subfamily Pomacentrinae: *Amphiprion akindynos*, *Chrysiptera flavipinnis*, *Dischistodus perspicillatus*, *Neoglyphidodon melas*, *Pomacentrus amboinensis*, *P. bankanensis*, *P. chrysurus*, *P. nagasakiensis*, and *P. pavo)* were collected from coral reefs around Lizard Island (14°40′S, 145°27′E), Northern Great Barrier Reef (GBR), between 2019 and 2022. Collections were conducted in the summer months (mid-November to early March) to avoid seasonal variability in body colouration and gene expression. Animal collection, husbandry and euthanasia followed procedures approved by The University of Queensland’s Animal Ethics Committee (2016/AE000304 & 2022/AE000569). The collections were conducted under permits from the Great Barrier Reef Marine Park Authority (G17/38160.2) and Queensland General Fisheries (207976).

We sampled three life stages per species: larval, juvenile, and adult specimens as, in a few species, significant morphological changes occur between these stages (e.g. *N. melas*)(Fig. S6-S16). Light traps were used to collect the fish at the end of their larval phase before settlement on the reef (Doherty, 1987). Light traps were placed 20-30 m from the reef at sunset and collected at dawn. Specimens were sorted in aerated tanks at the Lizard Island Research Station and processed the same morning. Juvenile and adult fish were collected on the reefs at depths of 1-15 m both on SCUBA and snorkel with hand nets and an anaesthetic clove oil solution (1/6 clove oil, 1/6 of 99% EtOH, and 4/6 of salt water), or using barrier nets. Fish were kept in flow-through aquaria at the research station, where they were exposed to natural sunlight for a maximum of 48 hours before further processing.

### Photography in RGB and UV

Three individuals per life stage per species (and n = 2 sub-adults of *D. perspicillatus*) were tentatively photographed in both the human visible (i.e., RGB) and the ultraviolet spectrum (we were able to collect only UV images of the larval stage for only six species) using two Nikon D810 cameras, one of which had its original filters removed to allow for full-spectrum sensitivity (Anderson Camera Repairs, Brisbane). Each camera was equipped with either a Nikon AF-S Nikkor 50 mm F/1.8G or a Nikon Micro-Nikkor 60 mm F/2.8 lens, depending on the size of the subject. The full-spectrum camera was constrained to ultraviolet wavelengths with a Schott UG11 visible-blocking filter and a Newport FSQ-BG39 blue bandpass filter. See the supplements (Figs. S3 and S4) for spectral sensitivity curves.

Each fish was placed in a small glass aquarium and allowed to acclimate for 2-3 min before being photographed by both cameras in sequence. Lighting was provided by two Nikon Speedlight SB-26 flashes placed at opposite corners of the aquarium, pointing down and inwards at the subject at 45°. Both speedlights were stripped of their filters and diffusers for full spectrum illumination (Fig. S5). Particularly for bigger fish, a large aperture was used to increase the depth of field and ensure the whole individual was in focus. Image J v1.53k was subsequently used to convert raw images to black and white. Processed images were then visually scored for the presence and type of UV patterns: i) no UV reflectance/colour present, ii) simple UV pattern defined as either uniform stripes or uniform UV body reflectance (see Fig. 1C), iii) spatially complex UV patterns defined as facial or body patterns with intricate, differently shaped dotted and lined motifs (see Fig. 1A and S11-S15, Table S10).

**Figure 1.**
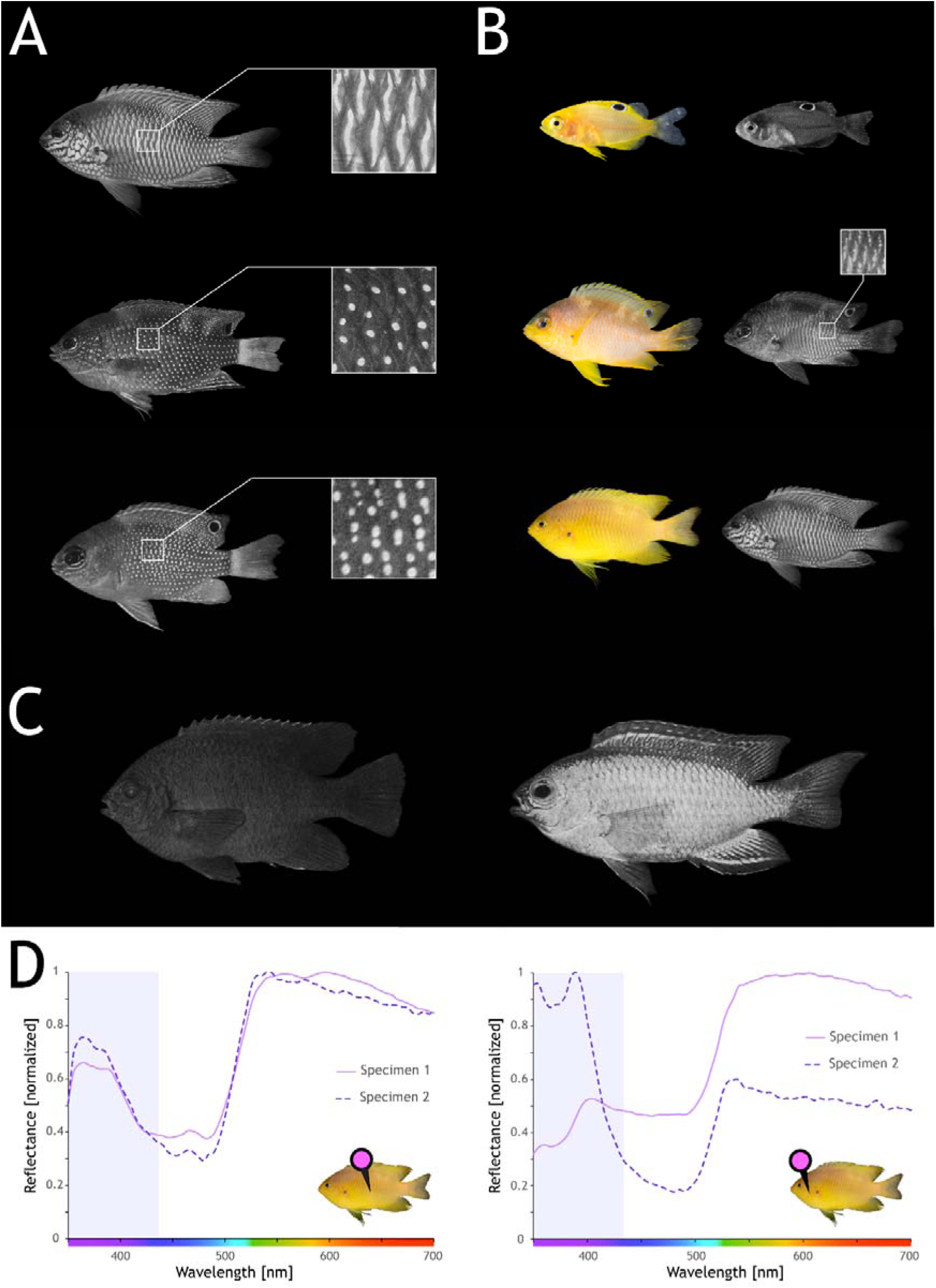
Ultraviolet colour (UV) pattern development and diversity in damselfishes. A) From top to bottom, diversity of complex UV patterns in the adults of *Pomacentrus amboinensis*, *P. bankanensis* and *P. chrysurus*. B) Development of colouration in the visible (left) and UV (right) spectrum in *P. amboinensis*. From top to bottom, the larval, the juvenile and the adult stage. The onset of the UV patterns is highlighted in the juvenile stage. C) Examples of other species visually scored for presence/absence and type of UV pattern. From left to right: *Neoglyphidodon melas* (no UV reflectance/patterns), and *Chrisyptera flavipinnis* (simple UV pattern). D) Spectral reflectance measurements on the body (left) and on the operculum (right) of *P. amboinensis* adults (n = 2). The violet box highlights the UV spectrum.

### Spectral reflectance measurements

To assess whether the damselfish UV patterns would differ in a spectral curve, i.e., if one individual uses different UV colours, we focused on *P. amboinensis*, for which complex UV (facial) patterns had previously been described (Siebeck, 2004; Siebeck et al., 2010) and were easily accessible. Spectral reflectance measurements of two adult *P. amboinensis* caught from the reefs around Lizard Island in 2005 and 2024 were obtained using an Ocean Optics (Dunedin, FL, USA) USB2000 spectrophotometer connected to a laptop computer running Ocean Insight (https://www.oceaninsight.com) OOIBASE32 or SpectraSuite software. Fishes were measured in the laboratory by removing them from the water and placing them on a wet towel to facilitate handling. Spectral reflectance curves measured this way do not significantly differ from those measured in water (Marshall et al., 2003). We used a PX-2 pulsed xenon light source (Ocean Insight) connected to a 400 μm UV-visible fibre, and the colour reflectance was measured using a 100 μm UV-visible fibre connected to the laptop.

The bare end of the collecting fibre was held at a 45° angle to prevent specular reflectance. A Spectralon 99% white reflectance standard was used to calibrate the percentage of light reflected at each wavelength from 350 – 750 nm. Measurements were taken from the operculum and the centre of the body, where different types of UV patterns occur (Siebeck et al., 2010; also see Fig. 1). Between 4 – 10 measurements per individual were taken from each location and subsequently averaged.

### RNA Extraction, Sequencing and Transcriptome Assembly

Fish were first photographed (see below) before being sampled for RNA sequencing (n = 3, samples per stage per species). Briefly, larvae were anaesthetised and killed between 7 am and 11 am and subsequently stored, as a whole, in RNAlater (Thermofisher) at –20 °C. Juveniles and adults were anaesthetised and killed by decapitation between 1 pm and 5 pm; retinas were removed from the eyecup and then stored in RNAlater at –20 °C.

Barcode DNA sequencing of the Cytochrome *c* oxidase I (COI) region was used to identify larvae that could not be assigned to a species based on morphology alone (n = 2 species). Briefly, DNA was extracted from fin clips using a custom-made lysis buffer (30 µl of 50 mM NaOH) and incubated at 90° C for 30 minutes. Subsequently, a neutralising buffer was added to the solution (5 µL Tris-HCl 1M pH 8.0). The samples were briefly vortexed and spun down; 2 µl of the supernatant was used to run a polymerase chain reaction (PCR) using fish universal *COI* primers (Ward et al., 2005) (for detailed methods, see Supplementary Methods). DNA was purified from the PCR product with the Monarch® PCR & DNA Cleanup Kit (New England Biolabs, https://www.nebiolabs.com.au/) and submitted for Sanger sequencing to the Australian Genomic Research Facility (https://www.agrf.org.au/). *COI* sequences were assessed using Geneious Prime v.2023.0.4 (https://www.geneious.com/), and low-quality bases were removed after visual inspection. Cleaned *COI* barcodes were then uploaded to the Barcode of Life Data System (http://v3.boldsystems.org/; Ratnasingham and Hebert, 2007) for species identification using default settings.

Larval eyes were removed from the eyecup using forceps and then homogenised using pestles (Interpath Services, https://www.interpath.com.au/). Juvenile and adult retinas were homogenised using glass beads (Sigma, https://www.sigmaaldrich.com/) and a TissueLyser LT (Qiagen, https://www.qiagen.com/). Following homogenisation, total RNA was extracted with the RNeasy Mini Kit (Qiagen) following the manufacturer’s protocol. An optional DNase step was performed to remove any trace DNA.

RNAseq library preparation using the NEBNext Ultra RNA library preparation kit for Illumina (NEB; mRNA Poly-A enriched, non-stranded library) and transcriptome sequencing were outsourced to Novogene (Singapore, https://www.novogene.com/). The concentration and quality of libraries were assessed by a Qubit dsDNA BR Assay kit (ThermoFisher) before barcoding and pooling at equimolar ratios. The libraries were then sequenced on a Hiseq2500 (PE150, 250–300 bp insert, ∼20 M fragments/library). Retinal transcriptomes were filtered and *de novo* assembled as per de Busserolles et al., 2017 and Tettamanti et al., 2019. Briefly, raw transcriptomes were uploaded to the Galaxy Australia server (https://usegalaxy.org.au/) (Afgan et al., 2018), filtered by quality using Trimmomatic (Galaxy version 0.36.6) (Bolger et al., 2014), then *de novo* assembled with Trinity (default settings, paired-end, --min_kmer_cov = 4; Galaxy version 2.9.1) (Haas et al., 2013).

### Opsin gene mining and phylogenetic reconstruction

To investigate developmental changes in damselfish fish vision in more detail, we mined the damselfish retinal transcriptomes for opsin gene sequences, following the protocols in de Busserolles et al., 2017 and Tettamanti et al., 2019. Briefly, we used the opsin gene coding sequences of the dusky dottyback, *Pseudochromis fuscus* (Cortesi et al., 2016), as a reference against which to map the assembled transcriptome of each individual in Geneious Prime. The *P. fuscus* opsin repertoire was chosen as a reference as the species is relatively closely related to the damselfishes, and it possesses all orthologues of the ancestral vertebrate opsin genes (Cortesi et al., 2016). Because lowly expressed genes are often overlooked in short-read assemblies and highly similar genes, such as the opsins, suffer from chimeric assembly errors, we also used a second approach to verify our initial findings, as per Musilova et al., 2019. Briefly, the raw transcriptome reads were mapped against the mined damselfish opsin gene sequences (fine-tuning, none; maximum gap per read, 10%; word length, 18; maximum mismatches per read, 2%; maximum gap size, 12 bp; and index word length, 14). Moving from single nucleotide polymorphism (SNP) to SNP, reads were manually extracted, taking advantage of the paired-end information between the SNPs. The extracted reads were then *de novo* assembled, and their consensus was extracted to form the entire coding region of the opsin gene. Opsin identity was verified through BLAST (https://blast.ncbi.nlm.nih.gov/) and by phylogenetic reconstruction with a reference dataset of vertebrate opsins obtained from GenBank (www.ncbi.nlm.nih.gov/genbank/) and Ensembl (www.ensembl.org).

The opsin gene phylogeny (Fig. S1) was obtained by first aligning the damselfish opsin genes to the reference dataset using the L-INS-I settings of the Geneious MAFFT plugin v1.5.0 (Katoh and Standley, 2013). The two previously described *sws1* copies of *A. ocellaris* (Mitchell et al., 2021) were added to the dataset to infer the identity of the orthologues in other species. jModeltest v2.1.6 (Darriba et al., 2012) was used to determine which model of sequence evolution was the most appropriate based on the Akaike information criterion. The phylogeny was then inferred using MrBayes v3.2.7a (Ronquist et al., 2012) as part of the CIPRES platform (Miller et al., 2010) using the following parameters: GTR+I+G model; two independent MCMC searches with four chains each; 10 M generations per run; 1000 generation sample frequency; and 25% burn-in.

The same settings were used to infer the phylogeny of the *sws1* gene clade. However, in this case, we only used the first and the fourth exons for the alignment, as they carried the strongest phylogenetic signal. Gene conversion between *sws1* copies had confounded the signal of the remaining exons (see below) (Fig. S2). After confirming the identity of the *sws1* copies, they were plotted against the most recent damselfish phylogeny (McCord et al., 2021) to reconstruct the evolutionary history of the gene in the Pomacentridae family.

### Sws1 gene conversion

We used GARD (Genetic Algorithm for Recombination Detection) (Kosakovsky Pond et al., 2006), with default settings to search for patterns of gene conversion between the *sws1* paralogs (n = 16, for nine Pomacentrinae species). Domains between breakpoints, i.e., the sections of putative sequence exchange, were subjected to phylogenetic comparisons to identify the most likely ancestry for each section.

### Sws1 spectral-sensitivity predictions

Amino acid comparisons at known tuning sites were used to estimate the spectral sensitivities of damselfish SWS1-based visual pigments following the methods in Mitchell et al., 2021 and Stieb et al., 2024. Briefly, SWS1 amino acid sequences of the eleven damselfish species were aligned to bovine rhodopsin (BRH; PBD accession no.1U19) as a reference. We focused on the SWS1 amino acids at BRH sites 114 and 118 as these sites have previously been shown to evolve in tandem and to confer peak spectral sensitivity shift of ∼10 nm (Stieb et al., 2024). SWS1 copies with BRH sites A114 and A118 were classified as shorter-wavelength sensitive (λ_max_ 356-362 nm), and copies with BRH S114 and S118 were classified as longer-wavelength sensitive (λ_max_ 368-370 nm), as per Stieb et al., 2024. These estimations assume a visual pigment with an A_1_-based chromophore. A_1_ is the dominant chromophore in coral reef fishes (Toyama et al., 2008), and based on the retinal transcriptomes, none of the species investigated in this study expressed *cyp27c1* (data not shown), the enzyme needed to convert A_1_-based chromophores to A_2_-based chromophores.

### Opsin gene expression and analysis

Opsin gene expression was calculated by mapping the filtered transcriptome reads for each individual to the species-specific opsin coding regions as per Tettamanti et al., 2019. The number of mapped reads (R) was normalised to the length (bp) of the opsin gene (i) to which they were mapped against:

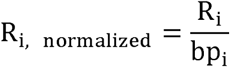

The proportional gene expression (p) for single (p_SC_) and double cone (p_DC_) opsins out of the total normalised expression for each cone type (T_SC_; T_DC_) was then calculated using the following equations:

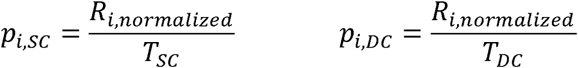

The proportional gene expression of the rod opsin (p_rod_) was calculated by comparing it to the total normalised opsin expression (T_opsin_):

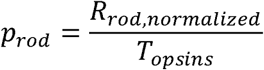

Expression plots were generated in Rstudio v1.4.1106 (Allaire, 2012), using a customised R script (R version 4.1.0, Ihaka and Gentleman, 1996).

### Differential gene expression

To investigate differences in gene expression throughout ontogeny, we first performed a Principal Component Analysis (PCA) using all samples (n = 93). Based on the resulting three clusters (Fig. 2A), we selected *P. amboinensis, Chro. atripectoralis* and *A. akindynos* as representative species for more in-depth analyses of the top differentially expressed genes (DEGs) between developmental stages. Due to the unavailability of a high-resolution genome for any of the species investigated, the filtered RNAseq reads were mapped to that of a close relative, the false percula anemonefish, *Amphiprion ocellaris* (subfamily Pomacentrinae; NCBI accession number: GCF_022539595.1) (Ryu et al., 2022) on the Galaxy Australia server. Mapping was performed using HISAT2 v2.2.1 with default parameters (Kim et al., 2019) to create a mapping-based estimation of transcript abundance. The function htseq-count v0.9.1 from HTSEQ (Anders et al., 2015) was used to quantify the number of mapped reads per gene based on the reference *A. ocellaris* genome annotation. The function generate-count-matrix v1.0 was then used to create a read count data table.

**Figure 2.**
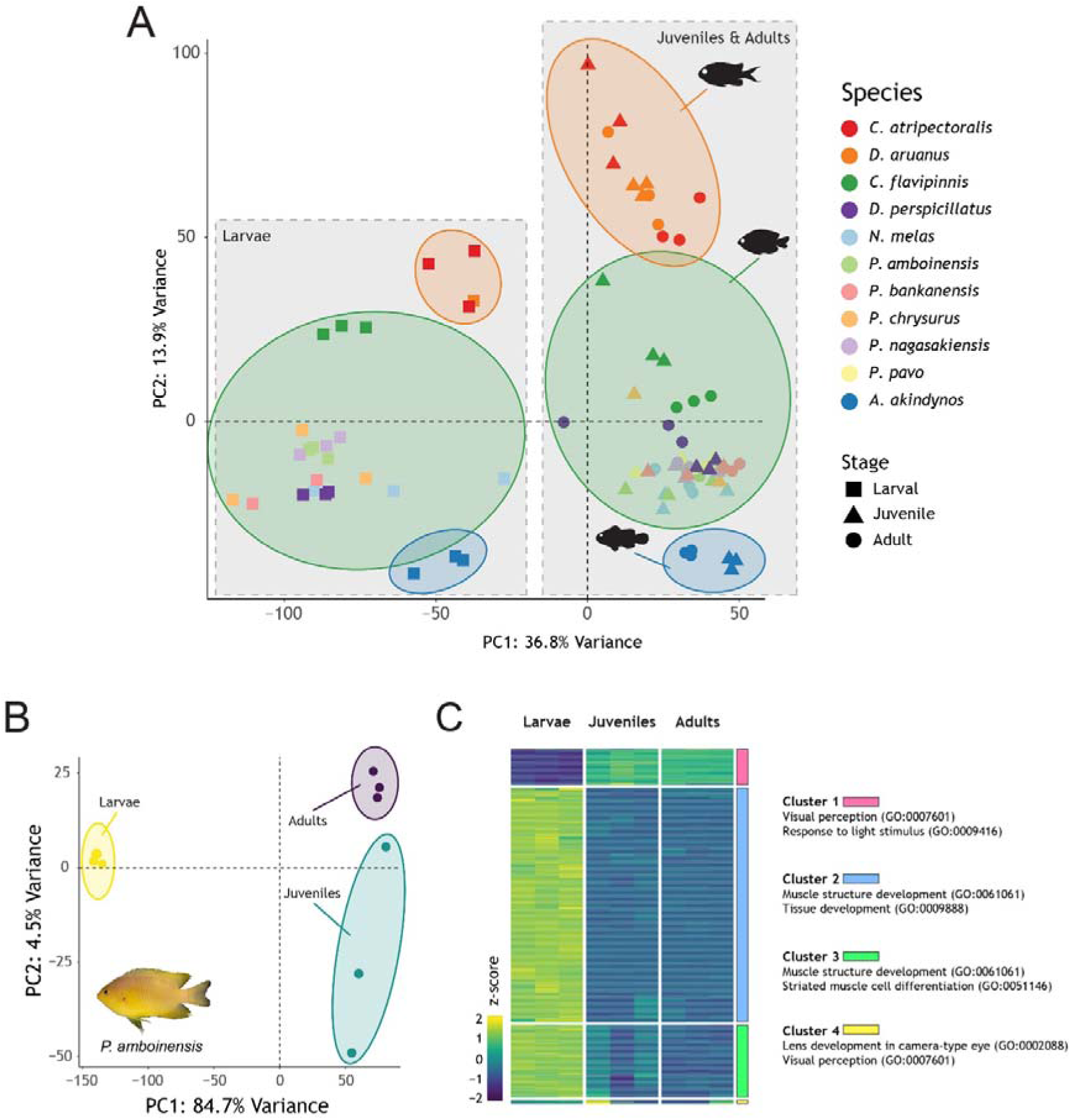
Differential gene expression analysis of damselfish retinal transcriptomes. A) PCA of the larval, juvenile, and adult stages of the eleven species investigated in this study (n = 93 samples). The two grey boxes indicate the major clusters derived from a split between the expression in larvae vs juveniles and adults. Three major species clusters are also portrayed. These are, from the top to the bottom: a cluster comprising the two species from the Chrominae sub-family (*Dascyllus aruanus* and *Chromis atripectoralis*); a second cluster comprising most Pomacentrinae species (all *Pomacentrus* spp., *Chrysiptera flavipinnis*, *Neoglyphidodon melas* and *Dischistodus perspicillatus)*; a third cluster for the anemonefish *Amphiprion akindynos*. B) PCA of the three developmental stages of *P. amboinensis*. Larval, juvenile, and adult samples cluster separately. C) Heatmap depicting scaled (z-score) expression levels of the top 1000 variable genes in the three stages of *P. amboinensis*. Genes are grouped on the y-axis into four major clusters based on their gene ontology (GO) enrichment analysis.

Differential gene expression between larval, juvenile and adult fish was inferred using DeSeq2 (Love et al., 2014), on the iDEP.96 platform (http://bioinformatics.sdstate.edu/idep96/; Ge et al., 2018). Briefly, the read count data was uploaded to iDEP.96 and matched automatically to the zebrafish genome assembly GRCz11 (*Danio rerio*; GCF_000002035.6), and gene symbols were converted to ENSEMBL gene IDs (https://asia.ensembl.org/index.html) for subsequent enrichment analysis. Pre-processing included removing features with less than 0.5 counts per million across all samples and transforming the data with the rlog algorithm from DESeq2. *P. amboinensis* filtered data was then used to perform a PCA and to create a heatmap of the top 1000 variable expressed genes, grouped into major clusters by k-means based on the elbow method to infer the optimal number of clusters (Fig. 2B, C). An enrichment pathway analysis of each cluster was then performed, and the top two GO annotations, sorted by False Discovery Rate (FDR), were selected to define the biological function of each cluster.

DESeq2 was used in the three species selected to find DEGs with a p-adjusted value of <=0.05 and a minimum fold-change (FC) of 2. Three comparisons were made: larval vs. juveniles, larval vs. adults, and juvenile vs. adults. The top 15 up- and down-regulated genes, as determined by log fold-change of each comparison, were used to perform a GO enrichment analysis in PANTHER via The Gene Ontology Resource (Tables S1-9) (Ashburner et al., 2000; Thomas et al., 2022). In the majority of cases, differentially expressed genes had Ensembl gene IDs allocated (i.e., LOC followed by the NCBI gene ID, e.g. LOC111574217), and corresponding gene orthologues from zebrafish or medaka (*Oryzias latipes*) were searched for using OrthoDB (Kuznetsov et al., 2023). If a GO function was missing from a gene, AMIGO was used to infer the function based on vertebrate orthologues (Carbon et al., 2009).

## Results

### Ontogenetic changes in UV colour and patterns

UV and human-visible photography showed that none of the species had UV patterns at the larval stage, and *N. melas* was the only species with no UV colouration as an adult. Simple UV patterns were discovered in juvenile and adult *A. akindynos*, *Chro. atripectoralis*, *Chry. Flavipinnis*, *D. aruanus*, and *N. melas* juveniles. We identified complex UV patterns with variable stripes, lines and dotted motifs in later ontogenetic stages of all *Pomacentrus* species (*P. amboinensis*, *P. bankanensis*, *P. chrysurus*, *P. nagasakiensis*, *P. pavo*), and *D. perspicillatus* (Fig. 1 for examples; Fig. S6-S16 for all species). In the *Pomacentrus* spp., the complex UV patterns first appeared at the juvenile stage, and they maintained the patterns after that, except for *P. chrysurus,* which did not show the patterns as adults (Table S10; Fig. S14). For *D. perspicillatus*, the complex UV patterns were initially only found in the adult stage. Further analysis of sub-adult individuals (fish that were judged to be in-between juveniles and adults based on intermediate sizes, being tolerated within the territories of mature individuals, and due to human-visible patterns in-between stages) revealed that the UV patterns first emerge at this stage (Fig. S10). Spectral reflectance measurements of the complex UV patterns on the body and operculum of two individuals of *P. amboinensis* revealed differences in the shape and the UV-peak of the reflectance curves, with peaks of the body being around ∼365nm and the facial patterns on the operculum having reflectance closer to ∼395nm (Fig. 1D).

### Opsin gene mining and phylogenetic reconstruction

Transcriptome mining and subsequent phylogenetic reconstruction revealed that all damselfishes expressed a rod opsin (*rh1*) and at least four cone opsins (*lws*, *rh2a*, *rh2b* and *sws1*) in their retinas (Fig. S1). Two species (*P. amboinensis* and *P. nagasakiensis*) expressed a second green-sensitive *rh2a* copy, and *Chry. flavipinnis* expressed a second red-sensitive *lws* gene. Several species also expressed the violet-sensitive *sws2b* gene. Six species (*A. akindynos*, *N. melas*, *P. amboinensis*, *P. bankanensis*, *P. chrysurus*, and *P. nagasakiensis*) expressed two UV-sensitive *sws1* copies.

The separate *sws1* phylogeny based on exons 1 and 4 revealed that the single *sws1* expressed in *D. auranus* and *C. atripectoralis* formed a sister clade to the *sws1*α and *sws1*β duplicates (Fig. 3). Also, *sws1* in *D. perspicillatus* and *C. flavipinnis* fell within the α clade; the two *sws1* paralogs in the remaining Pomacentrinae species could be assigned confidently to either clade.

**Figure 3.**
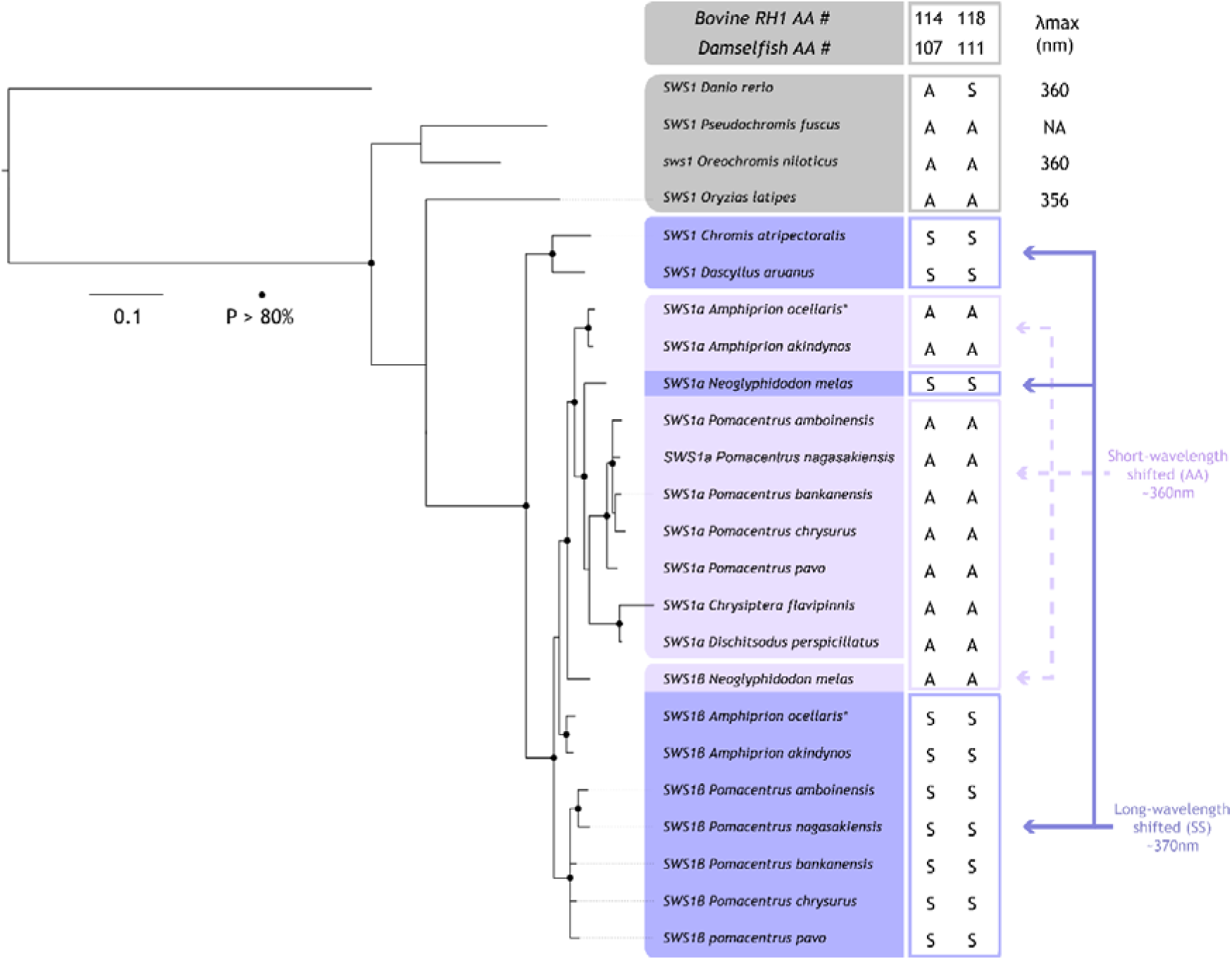
Evolution of *sws1* opsin genes in damselfishes, Pomacentridae. A) Bayesian phylogenetic tree of the relationship between *sws1* copies of eleven damselfish species. Annotated *sws1* copies of the false percula clownfish *Amphiprion ocellaris* (*) were used to infer the identity of the damselfish genes (Mitchell et al., 2021a). The amino acid (AA) identities at two key SWS1 spectral tuning sites, 114 and 118 (bovine rhodopsin numbering; Shi and Yokoyama, 2003; Yokoyama, 2000), are shown on the right. Each SWS1 opsin was classified as short- or long-wavelength sensitive based on the predictions of Stieb et al. 2024. In grey are SWS1 outgroup orthologs used for phylogenetic reconstruction showing the peak-spectral sensitivities (λ_max_) they convey. Note that, gene conversion between the *Neoglyphidodon melas sws1* copies has pulled *sws1*β towards the *sws1a* cluster and has led to an AA switch at the two tuning sites (also see Figs. 6 and S2 for details).

### Sws1 opsin evolution in damselfish

Plotting the *sws1* opsin genes onto the latest damselfish phylogeny (McCord et al., 2021) revealed that the *sws1* duplication most likely occurred in the Pomacentrinae ancestor (Fig. 3 and Fig. 4). The *sws1*β copy was most likely lost in the ancestor of the most ancestrally derived Pomacentrinae clade (Cheiloprionini as per Whitley, 1929; termed Pomacentrinae 1 in McCord et al., 2021; Tang et al., 2021), containing *D. perspicillatus* and *C. flavipinnis*.

**Figure 4.**
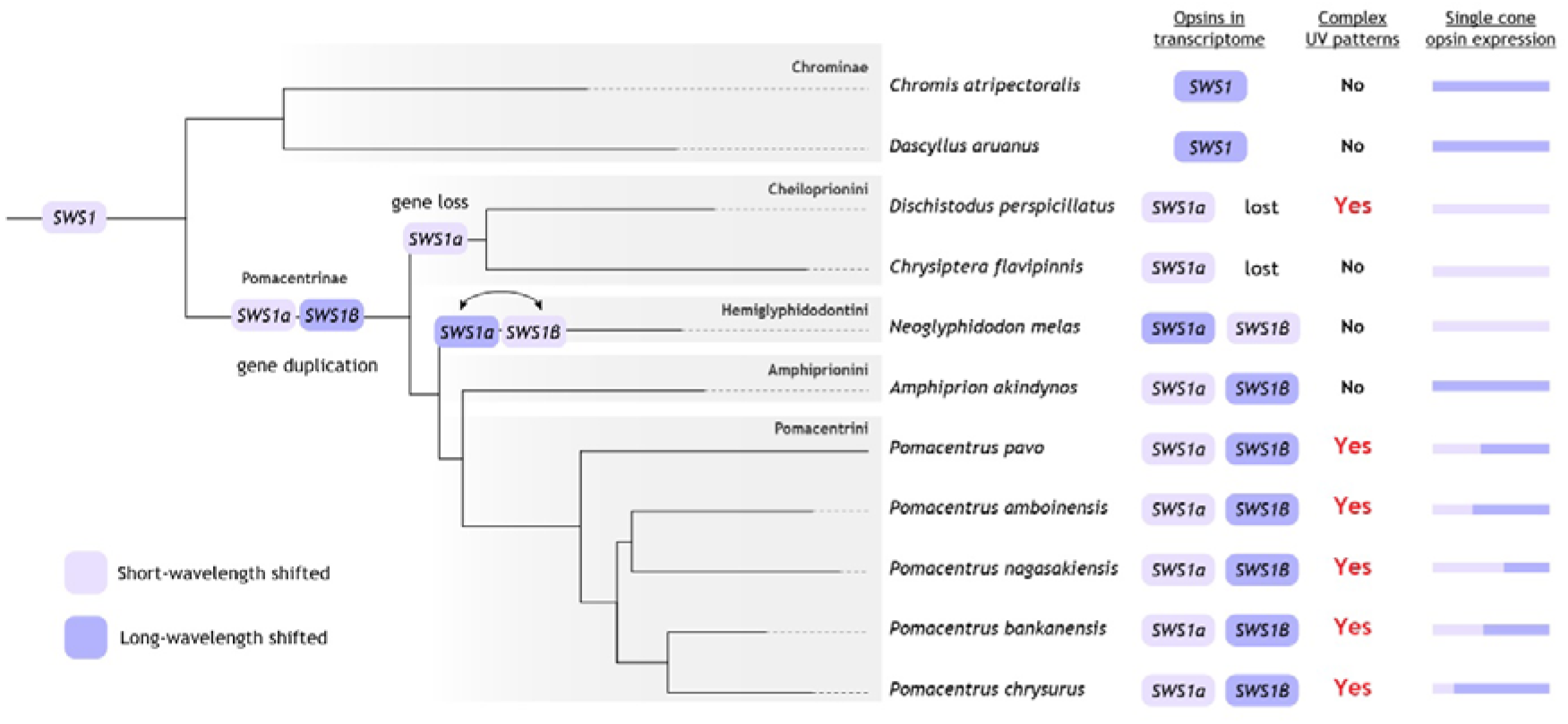
Overview of ultraviolet-(UV) wavelength sensitive *sws1* opsin gene evolution, gene expression and UV patterns in the eleven species of damselfish investigated. From the left: damselfish species tree, with the suggested timeline of gene duplication, loss, and gene conversion (double arrow); expression at any developmental stage of *sws1* in the retina; presence at any developmental stage of complex UV patterns; and relative *sws1* opsin expression in adult specimen (see Table 1 for detailed opsin gene expression data). Note that the *sws1* duplication likely occurred in the Pomacentrinae ancestor, and the *sws1*β copy was lost in the ancestor of the Cheiloprionini tribe. Gene conversion occurred between the *Neoglyphidodon melas sws1* copies (see Fig. S2 for details). The damselfish phylogeny was modified from McCord et al., 2021.

**Table 1.**
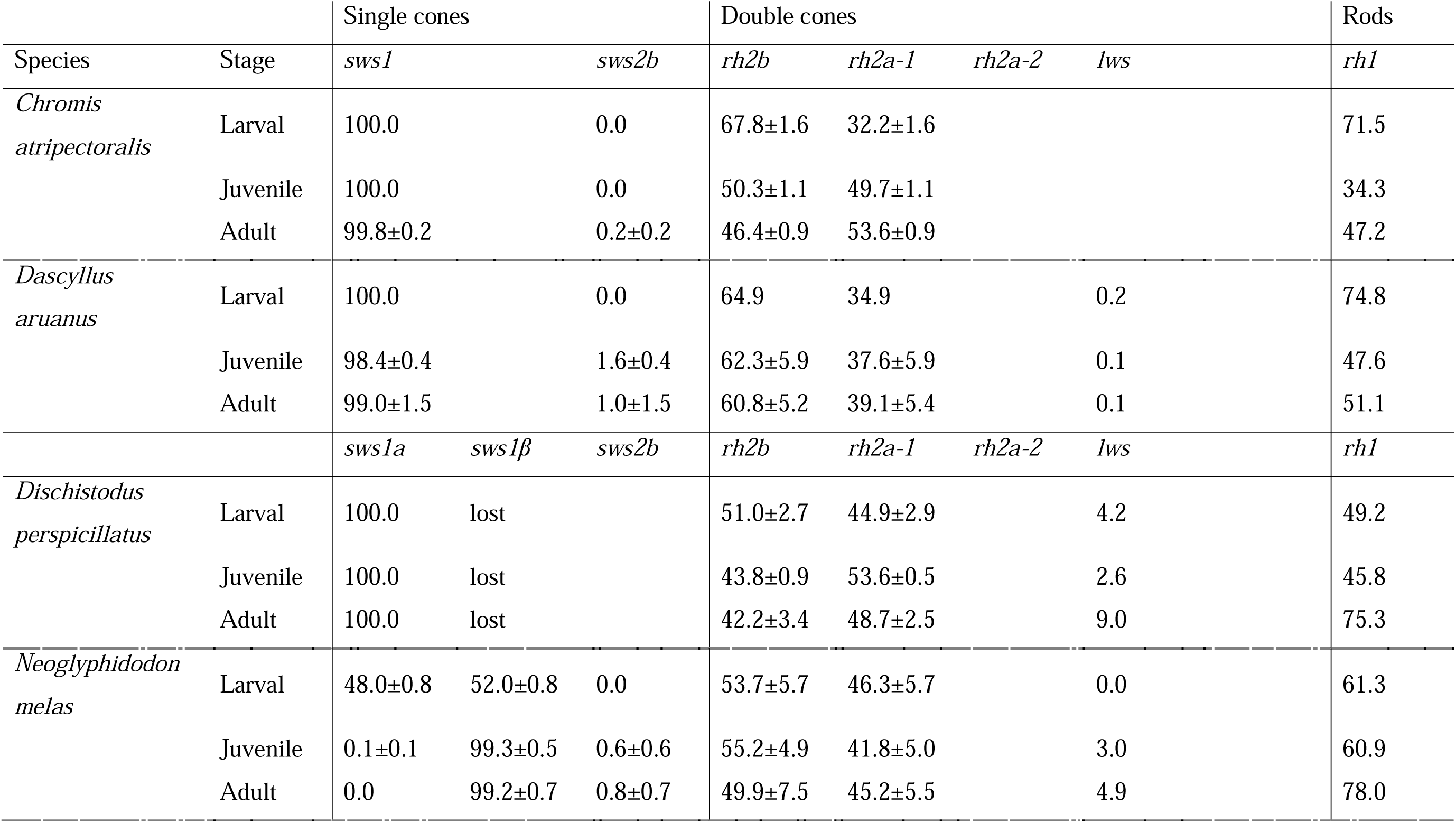
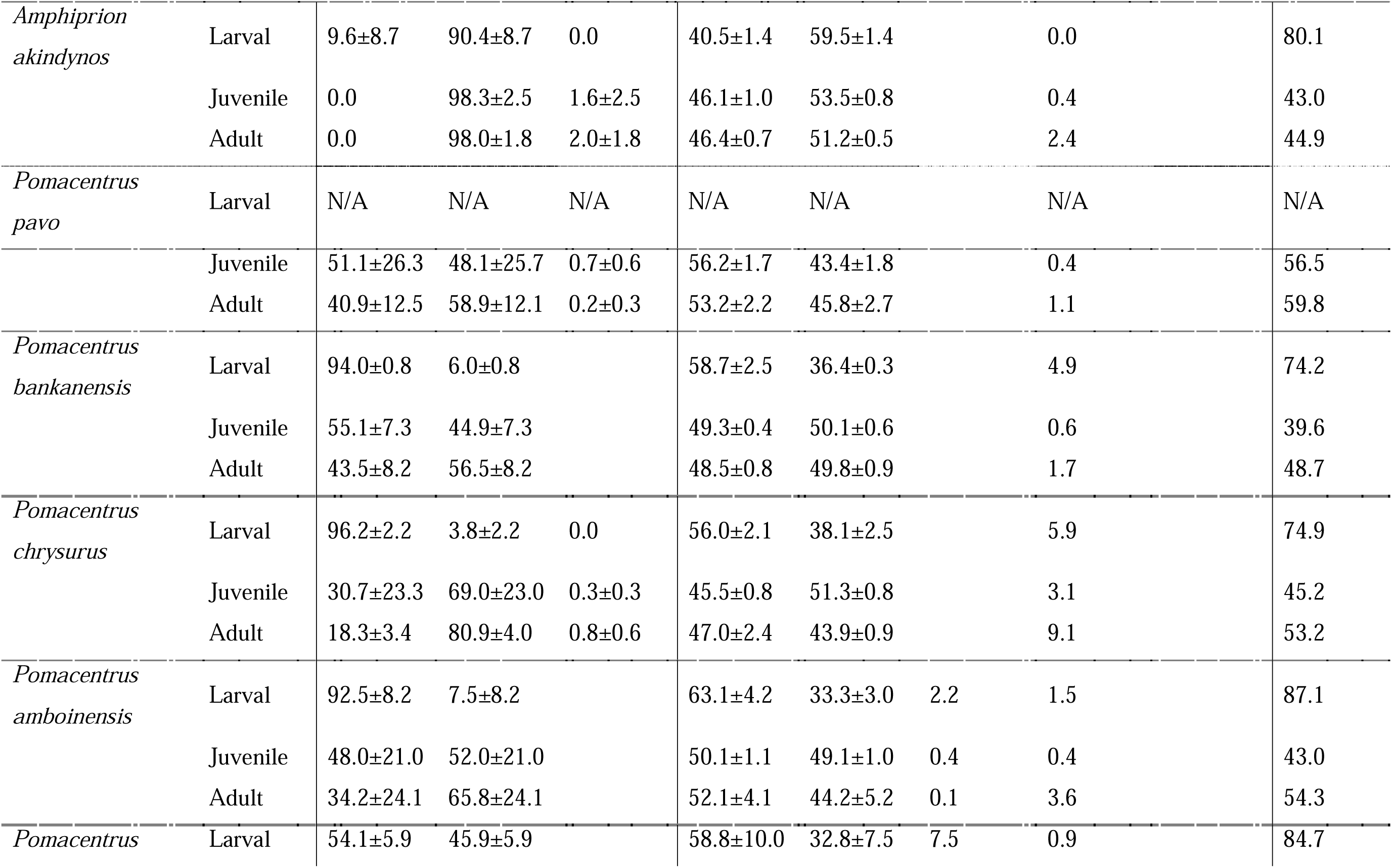
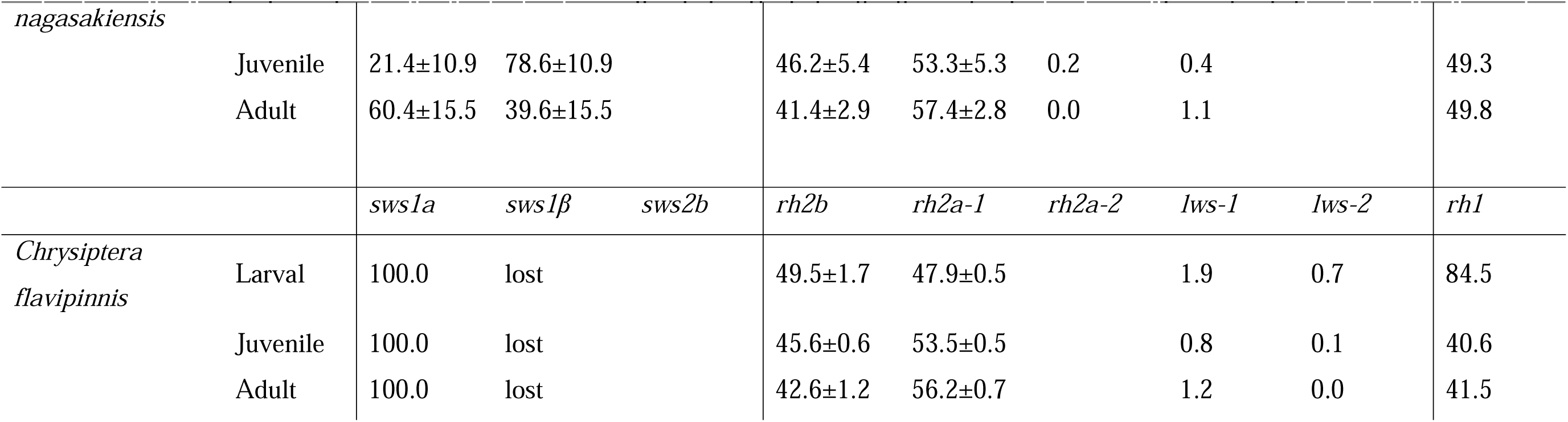
Opsin gene expression through development in eleven damselfish species. Shown are the proportional expression within single and double cones, and the rod-opsin expression versus total opsin expression.

Comparing the amino acids at the two major SWS1 tuning sites (BHR 114 and 118) revealed two distinct phenotypes: I) SWS1 orthologs with S114 and S118, predicted to be longer-wavelength shifted (∼370 nm λ_max_), II) SWS1 orthologs with A114 and A118, predicted to be shorter-wavelength shifted (∼360 nm λ_max_) (Fig. 3). SWS1 in the two Chrominae species, *D. auranus* and *C. atripectoralis*, and most SWS1β fall into group I, whereas most SWS1α belong to group II (Fig. 3).

Gene conversion analysis between the Pomacentrinae *sws1* paralogs revealed widespread conversion affecting different segments of the *sws1* genes, prompting us to rely only on exons 1 and 4 to resolve the *sws1* phylogeny (Fig. 3). The effects of gene conversion were most pronounced in *N. melas*, where *sws1*β was being pulled towards the *sws1a* clade, with the 114 and 118 tuning sites exchanged between copies (Fig. 3).

### Relative opsin gene expression through ontogeny

Using bulk retinal transcriptomes, we found that the relative opsin gene expression differed between species and ontogenetic stages (Table 1). Rod opsin, *rh1* expression was highest in larvae, except for *D. perspicillatus* and *N. melas*, in which the adults had the highest expression. Regarding the cone opsins, most species and stages predominantly expressed four to five genes (*sws1*α *&* β, *rh2b*, *rh2a*, and *lws*). However, *C. atripectoralis* expressed three cone opsin genes independent of life stage (*sws1*, *rh2b*, and *rh2a*), and *P. amboinensis* expressed six (*sws1*α *&* β, *rh2b*, *rh2a-1*, *rh2a-2*, and *lws*). There were notable differences in the expression of double-cone opsins with ontogeny. Generally, larval fish had a higher expression of *rh2b* than *rh2a*, with juveniles and adults exhibiting the opposite. *Lws* was lowly expressed or not expressed at all in most species at the larval stage. However, four species (*D. perspicillatus*, *C. flavipinnis*, *P. bankanensis*, and *P. chrysurus*) showed a higher expression (≥ 2% of proportional DC expression) at this stage. In the juvenile and adult stages, more species expressed *lws*. Notably, *N. melas*, *A*. *akindynos*, and *P. amboinensis* had > 3% proportional DC expression. For the SC, there were significant differences in the expression of *sws1* paralogs throughout ontogeny for six species (*P. amboinensis, P. nagasakiensis, P. chrysurus, P. bankanensis,* and *N. melas*), transitioning from a higher expression of *sws1*α in the larval stage towards a greater expression of *sws1*β in the later stages (Fig. 5).

**Figure 5.**
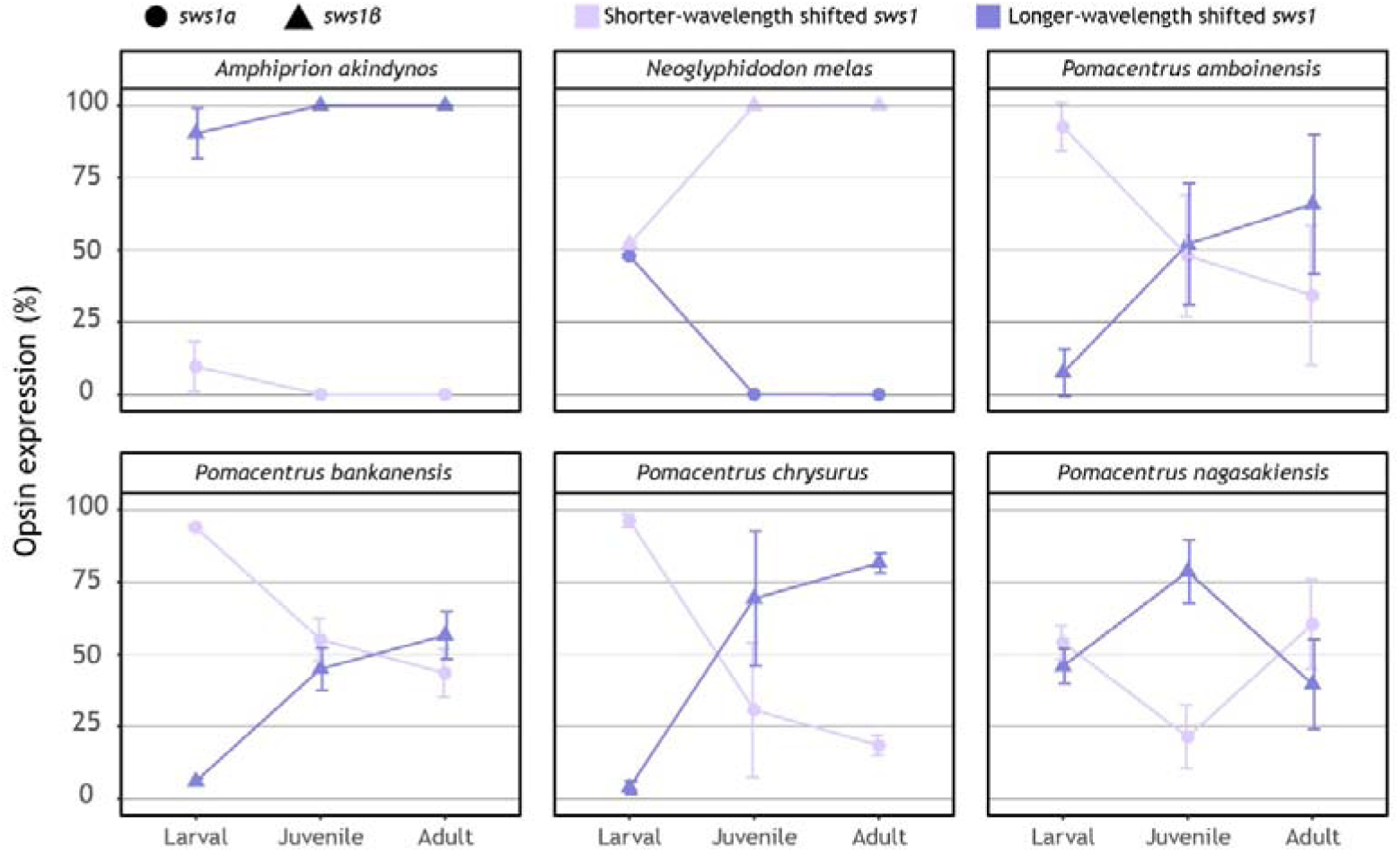
Ontogeny of *sws1* opsin gene expression in damselfish species with two paralogs. The two SWS1 copies are classified as short- and long-wavelength shifted based on Fig. 3. *A. akindynos* and *N. melas* transition from expressing both copies at the larval stage to only expressing *sws1*β at later stages. On the contrary, most *Pomacentrus* species primarily express *sws1*α at the larval stage, shifting towards a more balanced expression of the two *sws1s* in later stages.

### Differential gene expression

Retinal transcriptome assemblies and subsequent DGE mapping of all samples/species found 26,774 expressed transcripts, of which 11,227 passed the pre-processing filtering, and 7,769 transcripts were converted to Ensembl gene IDs (41.9% of total transcripts). PCA revealed that independent of species, larvae formed one cluster and juvenile and adult stages formed a second cluster along PC1, which accounted for 36.8% of the variance (Fig. 2A). Along PC2 (13.9% of the variance), three major clusters could be distinguished: one comprising the two Chrominae species (*Chro. atripectoralis*, *D. aruanus*), a second one including all the *Pomacentrus spp., Chry. flavipinnis*, *N. melas* and *D. perspicillatus*, and a third for the anemonefish *A. akindynos*.

An in-depth analysis of *P. amboinensis* separated the retinal gene expression profiles by life stage (n = 3 specimens/life stage; Fig. 2B). PC1 explained 84.7% of the variance, while PC2 explained 4.5%. A total of 23,190 transcripts were expressed in the retinas of the nine *P. amboinensis* samples, of which 20,952 passed the pre-processing filtering. Of these, 13,038 genes were converted to Ensembl IDs (56.2% of total transcripts). The top 1000 most variable genes clustered into four major groups based on their expression pattern across samples using k-means (Fig. 2C). GO enrichment analysis of overrepresented biological processes showed that three of the clusters were mostly comprised of genes upregulated in larvae, which were involved in developmental processes (muscle structure development [GO:0061061], tissue development [GO:0009888]). The fourth cluster comprised genes involved in visual processes (visual perception [GO:0007601], response to light stimulus [GO:0009416]) that were primarily upregulated in juvenile and adult *P. amboinensis* (Fig. 2C).

Details about the pairwise DGE analyses for the three life stages (larval vs. juvenile, larval vs. adult, juvenile vs. adult) in *P. amboinensis*, *C. atripectoralis* and *A. akindynos* are provided in Fig. 6 and Tables S1-9. In the larval vs. juvenile comparison, 4,325; 2,998, and 6,049 genes were upregulated in the larvae of each species, respectively. In contrast, 4,075; 1,830; and 4,175 were upregulated in the juveniles. In the larval vs. adult comparison, 4,591; 3,081; and 5,322 genes were upregulated in the larvae, and 4,121; 1,688; and 3,636 were upregulated in the adults. In the juvenile vs. adult comparison, 868, 811, and 630 genes were upregulated in the juveniles. In contrast, 435, 682, and 1,468 were upregulated in the adults. A GO enrichment analysis of the top 15 up- and down-regulated genes for each comparison revealed that most of the genes that were upregulated in the larval stage, when compared to later stages, were involved in developmental processes, predominantly of the lens (Fig. 6, Tables S1-9). Specifically, developmental genes upregulated in larvae included keratin 5 (*krt5*), involved in epidermal cell differentiation (GO:0009913); a crystallin, gamma M3 (*crygm3*), involved in lens development in camera-type eyes [GO:0002088]; and a periostin, osteoblast-specific factor b (*postnb*), involved in extracellular matrix organisation (GO:0030198). Conversely, genes upregulated in juveniles and adults, when compared to larval expression were primarily involved in visual processes and intracellular pathways including beta-carotene oxygenase 2b (*bco2b*), involved in retinal metabolic processes (GO:0042574); red-sensitive opsin-1 (*opn1lw1*), involved in visual perception (GO:0007601); and, the G protein-coupled receptor kinase (*grk1b*), involved in the cone photoresponse recovery (GO:0036368).

**Figure 6.**
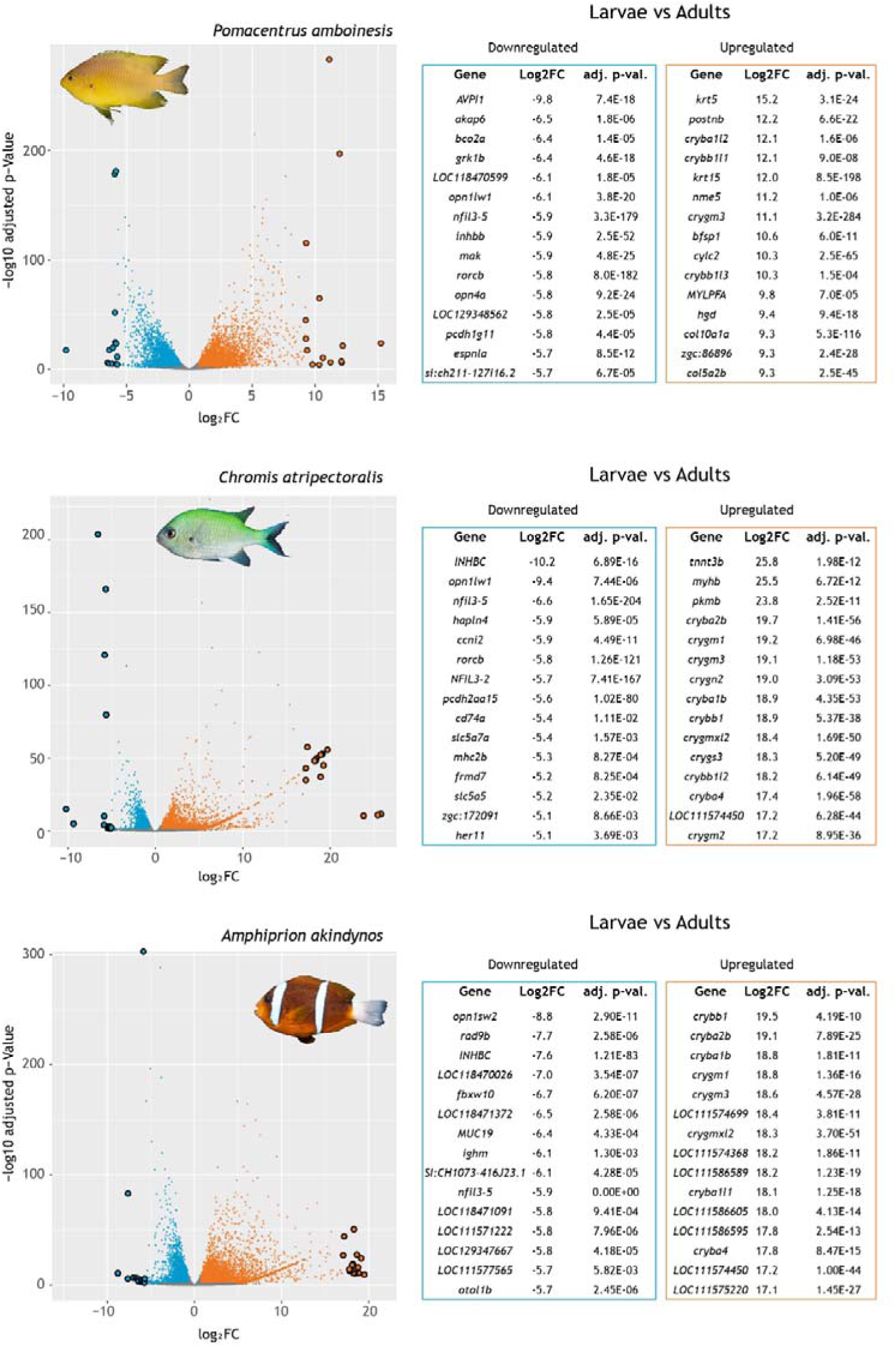
Differentially expressed retinal genes in larval vs. adult stages for three representative damselfish species (*Pomacentrus amboinensis*, *Chromis atripectoralis*, and *Amphiprion akindynos*), based on the three clusters from Figure 1A. Volcano plots depict retinal gene expression fold changes (log2FC) against their respective adjusted p-values (-log10). Genes that were significantly higher or lower expressed in larvae (adjusted p-value < 0.05, higher FC > 2, lower FC < -2) are represented in orange and blue, respectively. Grey dots represent genes that were not differentially expressed between stages. Enlarged outlined dots correspond to the 15 most up- or down-regulated genes for each comparison, with gene names in the respective tables. See Tables S1 – 9 for an extensive overview of the genes and their functions.

In the comparison between juveniles and adults, genes upregulated in the juvenile stage were involved in developmental processes and cellular division, such as the centromere protein W (*CENPW*), involved in the central nervous system development (GO:0007417); and the protein atonal homolog 7 (*atoh7*), involved in retinal development in camera-type eyes (GO:0060041). Genes upregulated in the adults were primarily involved in visual processes including relaxin family peptide receptor 3 (*rxfp3r*), involved in the G protein-coupled receptor signalling pathway (GO:0007186); cellular retinol-binding protein type II (*rbp2a*), involved in the vitamin A metabolic process (GO:0006776); and purpurin (*rbp4l*), which is involved in retinol transport (GO:0034633).

## Discussion

In this study we uncovered that all *Pomacentrus* species express two UV-sensitive *sws1* opsin genes at later developmental stages. The expression profile of these copies correlates with the appearance of complex UV patterns with variable stripes, lines and dotted motifs at the juvenile stage of *Pomacentrus* spp. In *P. amboinensis*, these complex patterns differ in spectral reflectance between pattern elements and might be used for individual recognition. We also found that larval fishes differ significantly in retinal gene expression and colouration from later stages, while differences between juveniles and adults were less pronounced. In the following we discuss each of our findings in detail.

### Ontogeny of damselfish UV colouration and visual adaptations

Using a comparative approach in eleven damselfish species, we show that complex UV facial and body patterns are more widespread than previously thought. Specifically, complex UV patterns were found in all *Pomacentrus* species and *D. perspicillatus* (Table S10, Fig. S10, S12-16). Previously, only the adults of the sister species of *P. amboinensis* and *P. moluccensis* have been shown to have complex facial UV patterns that they use for individual recognition (Parker et al., 2017; Siebeck, 2014, 2004; Siebeck et al., 2010). UV patterns were present from the juvenile stage onwards in *Pomacentrus* spp. (also see Gagliano et al., 2015 for previous work on *P. amboinensis*) and from the sub-adult stage onwards in *D. perspicillatus*. It is possible that complex UV patterns emerged a second time independently in *D. perspicillatus*. Alternatively, these patterns might have appeared in the Pomacentrinae ancestor and been lost multiple times after that (Fig. 4). Supporting the former hypothesis is that the patterns would have had to be lost at least four times independently (Pomacentrinae 1 – 4; McCord et al., 2021) in the Pomacentrinae history. The latter hypothesis is supported by the expression of two *sws1* copies correlating with complex UV patterns in *Pomacentrus* spp. Because the *sws1* duplication occurred in the Pomacentrinae ancestor, it is possible that the patterns emerged around the same time. This was then followed by a loss in complex patterns in species that retained only one *sws1* copy (e.g., *Chry. Flavipinnis*) or a simplification of patterns into stripes in others (e.g., *A. akindynos* and juvenile *N. melas*), with *D. perspicillatus* maintaining the ancestral trait but shifting its appearance to later in development (Fig. 4).

The expression of two or more *sws1* paralogues has so far only been reported from butterflies, snakes, mantis shrimp, a species of gar, and some adult damselfishes (mostly anemonefishes) (Bok et al., 2018; Briscoe et al., 2010; Cronin et al., 1994; Hauzman et al., 2021; Mitchell et al., 2021; Stieb et al., 2024; Sukeena et al., 2016). Here, we show a co-expression of the paralogs in *Pomacentrus* spp. In butterflies, the expression of two *sws1* copies allows for colour vision in the ultraviolet, with females able to discriminate between narrow UV spectra of 380 nm versus 390 nm (Finkbeiner and Briscoe, 2021). Similarly, mantis shrimp have been shown to distinguish between different UV colours (Bok et al., 2014). The expression of two *sws1* opsins that potentially confer different spectral sensitivities in the UV (∼15 nm apart) and the occurrence of differently coloured UV patterns in the fish (Fig. 1D) strongly suggest that *Pomacentrus* species are also able to distinguish between UV colours. However, in-situ hybridisation, single-cell RNA sequencing and microspectrophotometry are needed to show that the two damselfish *sws1* copies are in separate cone photoreceptors with different spectral sensitivities and their distribution across the retina. Finally, behavioural assays, like the ones in *A. ocellaris* (Mitchell et al., 2024), are necessary to prove UV-colour discrimination in these fishes.

### Sws1 evolution and spectral tuning

The phylogenetic reconstruction showed that the damselfish visual opsins mined from the retinal transcriptomes belong to the major five vertebrate clades (Yokoyama, 2000) (Fig. S1). A more detailed evolutionary reconstruction of the *sws1* genes revealed that the damselfish ancestor most likely possessed a single *sws1* copy (Fig. 4). These results are congruent with a more extensive phylogenetic comparison of damselfish *sws1* opsins conducted in Stieb et al., 2024. Also, by adding the transcriptomes of *D. perspicillatus* and *C. flavipinnis* (both Cheiloprionini), we could place the duplication of *sws1* at the base of the Pomacentrinae subfamily. Previously, the duplication was thought to have occurred only after the split of the Cheiloprionini from the rest of the Pomacentrinae (Stieb et al., 2024). Since only *sws1a* was recovered from the two Cheiloprionini species analysed here, and a single *sws1* was also recovered in the two *Chrysiptera* species from Stieb et al., 2024, it is likely that the tribe has lost *sws1*β ancestrally (Fig. 4). *N. melas* was the most ancestrally derived Pomacentrinae species from our dataset, with two copies (Fig. 4).

Interestingly, the *N. melas sws1* paralogs showed extensive patterns of gene conversion, with distinct breaking sites revealed by GARD analysis (Fig. S2) and *sws1*β being pulled towards the *sws1a* clade in the phylogeny (Fig. 3). This was also evident from the amino acid comparison at sites 114 and 118 (relative to BRH). Most damselfish *sws1*α orthologues had A114 and A118 (shorter-shifted phenotype), and *sws1*β had S114 and S118 (longer-shifted phenotype). However, the *N. melas* copies had the opposite amino acids at these two sites, with *sws1*α S114, S118 and *sws1*β A114, A118. GARD also revealed that gene conversion generally affected the *sws1* paralogs of Pomacentrinae, with higher sequence similarities detected in the first and last sections of the coding regions (Fig. S2). *N. melas* differed from the rest of the Pomacentrinae species in that the middle part, containing sites 114 and 118, was also affected. Future work should sample more broadly within the Hemiglyphidodontini (the tribe containing *Hemiglyphidodon, Amblyglyphidodon, Altrichthys, Acanthochromis* and *Neoglyphidodon;* Tang et al., 2021; Whitley, 1929) to reveal whether the pattern observed here is specific to *N. melas* or common to the tribe.

### Opsin gene expression

We found that damselfishes use a variety of visual opsin gene repertories with ontogenetic differences in gene expression. Consistent with previous research (Luehrmann et al., 2018; Stieb et al., 2024, 2023, 2019, 2017, 2016), each species expressed *rh1* and at least three cone opsin genes (*sws1*, *rh2a* and *rh2b*), with some species also expressing *sws2b*, *lws* and/or multiple *sws1*, *rh2a* and *lws* paralogs (Table 1). Some of this variation might be explained by short-term reversible changes in opsin gene expression, which are common in teleosts (reviewed in Carleton et al., 2020; Musilova et al., 2021) and have previously been documented in the adults of *P. amboinensis* and *P. moluccensis* (Luehrmann et al., 2018). Moreover, damselfishes are known to tune their visual pigments over evolutionary timeframes through changes in opsin gene sequences (Stieb et al., 2017). These adaptations are thought to be driven by species-specific ecologies. For example, some damselfishes may show intraspecific differences in *sws1* (UV-sensitivity) expression with depth (Stieb et al., 2016), and, as mentioned above, they might also show differences in *lws* (red-sensitivity) expression and spectral tuning that correlate with feeding mode and fish colouration (Stieb et al., 2023). As we show here, we also find consistent developmental plasticity in opsin gene expression between larval, juvenile and adult stages of all species (Table 1, Fig 4).

In damselfishes, such as *A. akindynos* (Stieb et al., 2019), *rh2a* and *rh2b* are expressed in opposite double cone members, forming a pair of visual pigments that is best tuned to the underwater light environment (blue-green) (Jerlov, 1976; Yokoyama and Jia, 2020). A slight change in *rh2b* gene expression with development was found for most species, which shift from a higher expression of *rh2b* in the settlement larval stage to a more even expression between *rh2b* and *rh2a* at later stages (Table 1). This could be explained by the open ocean light environment being blue-shifted compared to the broad light spectrum found on shallow reefs (all juveniles and adults, except for *C. flavipinnis*, were caught <10 meters in depth) (Jerlov, 1976; Marshall et al., 2003). This is similar to the intraspecific depth differences discovered previously in the adults of some damselfishes. Individuals caught at shallower depths had a higher expression of *rh2a* than *rh2b*, while a more even expression between genes was seen in deeper-caught specimens (Stieb et al., 2016). Changing opsin gene expression this way likely tunes the double cone spectral sensitivities to the most abundant wavelengths of light available in each life stage. However, as is the case in Killifish (*Lucania goodei*) (Fuller et al., 2004), this might result from environmental plasticity rather than a pre-determined part of development.

Two species (*P. amboinensis* and *P. nagasakiensis*) expressed a second *rh2a* paralog (*rh2a-2*). Interestingly, some, but not all, anemonefish species have also been found to have a second *rh2a* gene in their genomes (Mitchell et al., 2021; Musilova et al., 2019). The two damselfish species expressing *rh2a-2* diverged ∼20 Mya, while more closely related species, such as *P. chrysurus* (divergence from *P. nagasakiensis* ∼2.5 Mya; McCord et al., 2021), expressed only one *rh2a* copy. It is possible that the second copy was not expressed in most of the species analysed here. However, because of the abundant depth at which we sequenced the transcriptomes, we can usually detect traces of all visual opsins present in a species’ genome. Also, many more percomorph fishes have two *rh2a* gene copies (Lin et al., 2017; Musilova and Cortesi, 2023). This suggests that *rh2a* has duplicated early in percomorph evolution and that the orthologs were lost multiple times in different species. Unfortunately, extensive and frequent gene conversion between *rh2* genes has made it nearly impossible to conclusively disentangle their evolutionary history (Musilova and Cortesi, 2023).

S*ws1* was expressed in all stages and species investigated. Interestingly, species in the *Pomacentrus* clade, plus the larval stage of *N. melas*, were found to express two copies of *sws1*, which could be assigned to short- and long-wavelength shifted groups (see Fig. 3 for classification and Fig. 5 for expression). Multiple *sws1* genes are rare in fishes (Musilova et al., 2019) and in coral reef species have so far only been found in the damselfishes (Mitchell et al., 2021; Musilova et al., 2019; Stieb et al., 2024). The expression of *sws1* is often associated with UV communication and could also be an adaptation for feeding on UV-absorbing or reflecting zooplankton (Flamarique, 2016; Job and Bellwood, 2007; Novales-Flamarique and Hawryshyn, 1994; Siebeck, 2014, 2004; Siebeck et al., 2010; Stieb et al., 2017). Expression of *sws1* for foraging could be substantiated by damselfishes being highly opportunistic feeders; while pomacentrid species are often grouped into three major dietary classes (planktivore, herbivore and omnivore), they can readily shift feeding mode to exploit nutrient-rich food (Frédérich et al., 2006). Thus, retaining *sws1* expression even in prevalently benthic feeding species could benefit opportunistic predation on zooplankton and other small organisms. Feeding ecology, however, does not explain the expression of a second *sws1* gene in the *Pomacentrus* spp.

### Retinal gene expression throughout ontogeny

PCA revealed that in damselfishes, the retinal gene expression in larvae is distinctly different to later stages (Fig. 2A), suggesting that juvenile damselfishes are already displaying adult expression profiles. PCA and k-means clustering in *P. amboinensis* supported these findings by also showing a separation of gene expression between the larval and later stages (Fig. 2B, 2C).

Using DGE between developmental stages in three damselfish species (Fig. 6), the top 15 upregulated genes in larvae compared to juveniles and adults, independent of species, were primarily involved in developmental processes of the eye and especially in lens formation (e.g., *krt5*, *crygm3*, *postnb*) (Tables S1, S2, S4, S5, S8, and S9). This is similar to the retinal gene expression in settling surgeonfish larvae (Fogg et al., 2022b), and supports the metamorphosis seen in other organs of coral reef fishes during settlement on the reef (Hu et al., 2019; Lim and Mukai, 2014). Notably, various crystallin paralogues were upregulated in the larvae. These encode water-soluble proteins responsible for the transparency and the gradient of refractiveness of the vertebrate lens (Palczewski et al., 2000). The latter is a crucial aspect of teleost fish multifocal lenses, which enable focusing different wavelengths of light on the same plane (Gagnon et al., 2012; Karpestam et al., 2007; Kröger et al., 1999).

Compared to the larval stage, the top 15 upregulated genes in the juvenile and adult stages were primarily involved in visual processing and homeostasis of the retina (e.g., *grk1b*, *nfil3-5*) (Fig. 6). Specifically, an orthologue of the zebrafish beta-carotene oxygenase 2b (*bco2b*) was upregulated in juveniles and adults of *P. amboinensis*. In teleost fishes, beta-carotene oxygenases have evolved an extended repertoire through repeated gene duplications, and *bco2* paralogues are thought to participate in retinoic acid (RA) biosynthesis (Helgeland et al., 2014; Kiefer et al., 2001). Retinoic acid is involved in cone photoreceptor survival in the mouse retina (Amamoto et al., 2022). The upregulation of *bco2* in the juvenile and adult stages highlights the need for a continuous supply of RA for the long-term survival of their cones, which might be less critical in the still rapidly developing retina at the larval stage.

Unsurprisingly, the juvenile versus adult comparison also revealed an upregulation of developmental genes in the juvenile stage. For example, the atonal homolog 7 (*atoh7*), which is crucial for developing ganglion cells (Brown et al., 2001), was upregulated in juveniles compared to adults. Genes upregulated in adults compared to juveniles were primarily involved in visual processing and phototransduction, including the long-wavelength-sensitive opsin-1 (*opn1lw1* or *lws*). An increased proportional expression of *lws* was also present in the adults of several other damselfish species, including *D. perspicillatus*, *N. melas*, *A. akindynos*, and *P. chrysurus* (Table 1). This increase might be linked to changes in their feeding ecologies to a more herbivorous diet, as chlorophyll reflects strongly in the far-red, or due to the emergence of orange/red colours in later stages, as discussed in detail in Stieb et al., 2024.

## Acknowledgements

We would like to thank the Lizard Island Research Station staff for their support during fieldwork and acknowledge the Dingaal, Ngurrumungu and Thanhil peoples as traditional owners of the lands and waters of the Lizard Island region. We would also like to thank Professor Mark McCormick and his team for generously lending us their light traps and assisting with some animal collections and Dr Sam B. Powell for technical support with the UV camera system. We would like to acknowledge the Australian Museum and the Lizard Island Reef Research Foundation for supporting the fieldwork component of this research with a doctoral fellowship awarded to V.T. This work was supported by an Australian Research Council (ARC) Discovery Project (DP180102363) to J.M. and F.C., the AFOSR/AOARD to J.M., a UQ Amplify and ARC DECRA Fellowships (DE200100620) to F.C., and the Zoltan Florian Marine Biology Fellowship from the Lizard Island Reef Research Foundation (LIRRF) awarded to V.T.

## Data availability

Raw RNA sequencing data was uploaded to SRA (XXXX) and opsin genes sequences were uploaded to NCBI (XXXX) and will be made available upon acceptance. Transcriptome assemblies, pictures and phylogenetic trees are available on Dryad (XXXX).

## Author contributions

V.T. conceived the study and designed the experiments together with F.C., N.J.M. and K.L.C. V.T. performed molecular experiments and analysed gene expression data. V.T., F.C. and J.M. collected ultraviolet pictures and spectral reflectance from specimens. V.T. and F.C. wrote the initial manuscript and all authors contributed to the final submitted version.

