## Supporting_information for "Damsels in a hidden colour: development of ultraviolet sensitivity and colour patterns in damselfishes (Pomacentridae)"

|  | Gene ID | Gene name | logFC | Function |
| --- | --- | --- | --- | --- |
| Larvae | krt5 | Keratin 5 | 13.60953767 | epidermal cell differentiation |
|  | crygm3* | crystallin, gamma M3 | 11.90454196 | lens development in camera-type eye |
|  | krt15† | Keratin 15 | 10.53985011 | intermediate filament cytoskeleton organization |
|  | postnb | Periostin, osteoblast-specific factor b | 9.987486741 | extracellular matrix organization |
|  | cylc2* | Cylicin II | 9.710383086 | NA |
|  | nme5† | NME/NM23 family member 5 | 9.684379909 | cilium movement |
|  | ACTN3B | Actinin alpha 3b | 9.337753935 | actin cytoskeleton organization |
|  | COL5A2B | Collagen, type V, alpha 2b | 8.913992079 | extracellular matrix organization |
|  | cdh26.1 | Cadherin 26, tandem duplicate 1 | 8.771086861 | cell adhesion |
|  | mybpc2b | Myosin-binding protein Cb | 8.769956823 | muscle tissue development |
|  | zgc:86896 | Actin-related protein 2/3 complex subunit | 8.598860398 | regulation of actin filament polymerization |
|  | actc1a* | Actin alpha 1, skeletal muscle a | 8.508641207 | skeletal muscle fiber development |
|  | col1a1b | Collagen, type I, alpha 1b | 8.505589455 | extracellular matrix organization |
|  | myom2a | Myomesin 2a | 8.501334112 | extraocular skeletal muscle development^§^ |
|  | tp63 | Tumor protein 63 (p63) | 8.483962241 | central nervous system development |
| Juvenile | AVPI1 | Arginine vasopressin-induced protein 1 | -10.47233971 | cell cycle |
|  | bco2b* | Beta-carotene oxygenase 2b | -7.151599401 | retinal metabolic process |
|  | hmcn2* | Hemicentin 2 | -7.059654637 | extracellular matrix organization |
|  | parp6* | Poly (ADP-ribose) polymerase family, member 6a | -6.799079065 | endoplasmic reticulum unfolded protein response |
|  | akap6* | A kinase (PRKA) anchor protein 6 | -6.766419112 | positive regulation of potassium ion transmembrane transport |
|  | LOC129348730† | Ig kappa chain V region K16-167-like | -6.679892785 | immune response |
|  | inhbb* | Activin beta-B chain | -6.670850779 | SMAD protein signal transduction |
|  | mak* | Male germ cell-associated kinase | -6.652650352 | cilium assembly |
|  | espnla* | Espin-like a | -6.648552576 | actin filament bundle assembly |
|  | LOC129348562 | Uncharacterized loci | -6.439594702 | NA |
|  | si:ch211-127i16.2 | Amine oxidase | -6.388879033 | NA |
|  | LOC111576907 | Uncharacterized loci | -6.03194659 | NA |
|  | psme4b* | Proteasome activator complex subunit 4B | -6.031205638 | DNA repair |
|  | nfil3-5 | Nuclear factor, interleukin 3-regulated, member 5 | -5.915268133 | circadian rhythm |
|  | rorcb* | RAR-related orphan receptor C b | -5.88243781 | regulation of transcription by RNA polymerase II |
| *these genes were not properly annotated and thus orthologues from zebrafish (Danio rerio) or medaka (Oryzias latipes) were used in Gene ID for GO Enrichement analysis  †orthologues of these genes were not found, similar vertebrates genes found through BLAST (same PANTHER class) were used to infer the function  § GO function was missing and was inferred from other vertebrates orthologues | | | | |

**Table S1** Top 15 up- and down- regulated genes in the comparison of differential gene expressed between the larval and juvenile stage retinas of *P. amboinensis*.

|  | Gene ID | Gene name | logFC | Function |
| --- | --- | --- | --- | --- |
| Larvae | krt5 | Keratin 5 | 15.23444271 | epidermal cell differentiation |
|  | postnb | Periostin, osteoblast-specific factor b | 12.19589633 | extracellular matrix organization |
|  | cryba1l2 | BetaA1c-crystallin | 12.11578287 | lens development in camera-type eye |
|  | crybb1l1 | Crystallin, beta B1,-like 1 | 12.0989837 | lens development in camera-type eye |
|  | krt15 | Keratin 15 | 11.95142114 | intermediate filament cytoskeleton organization |
|  | nme5† | NME/NM23 family member 5 | 11.22791919 | cilium movement |
|  | crygm3* | Crystallin, gamma M3 | 11.13005248 | lens development in camera-type eye |
|  | bfsp1† | Filensin | 10.62476355 | lens development in camera-type eye |
|  | cylc2* | Cylicin II | 10.34511181 | NA |
|  | crybb1l3 | Crystallin, beta B1,-like 3 | 10.33116294 | lens development in camera-type eye |
|  | MYLPFA | Mylz2 protein | 9.796417856 | skeletal muscle tissue development |
|  | hgd | Homogentisate 1,2-dioxygenase | 9.374225136 | alpha-amino acid metabolic process |
|  | col10a1a* | Collagen, type X, alpha 1a | 9.296577307 | extracellular matrix organization |
|  | zgc:86896 | Actin-related protein 2/3 complex subunit | 9.277282415 | regulation of actin filament polymerization |
|  | col5a2b | Collagen, type V, alpha 2b | 9.259857629 | extracellular matrix organization |
| Adult | AVPI1 | Arginine vasopressin-induced protein 1 | -9.801142964 | cell cycle |
|  | akap6* | A kinase (PRKA) anchor protein 6 | -6.506175909 | positive regulation of potassium ion transmembrane transport |
|  | bco2a* | Beta-carotene oxygenase 2a | -6.383776877 | retinal metabolic process |
|  | grk1b | G protein-coupled receptor kinase | -6.365512676 | cone photoresponse recovery |
|  | LOC118470599 | Uncharacterized loci | -6.138620969 | NA |
|  | opn1lw1* | Red-sensitive opsin-1 | -6.079753307 | visual perception |
|  | nfil3-5 | Nuclear factor, interleukin 3-regulated, member 5 | -5.934511407 | circadian rhythm |
|  | inhbb* | Activin beta-B chain | -5.919933954 | SMAD protein signal transduction |
|  | mak* | Male germ cell-associated kinase | -5.88860943 | cilium assembly |
|  | rorcb* | RAR-related orphan receptor C b | -5.837080605 | regulation of transcription by RNA polymerase II |
|  | opn4a | Melanopsin-A | -5.826500894 | cellular response to light stimulus |
|  | LOC129348562 | Uncharacterized loci | -5.825187759 | NA |
|  | pcdh1g11* | Protocadherin gamma-A11 | -5.764863403 | nervous system development |
|  | espnla | Espin-like a | -5.740183945 | actin filament bundle assembly |
|  | si:ch211-127i16.2* | Amine oxidase | -5.728239063 | NA |
| *these genes were not properly annotated and thus orthologues from zebrafish (Danio rerio) or medaka (Oryzias latipes) were used in Gene ID for GO Enrichement analysis  †orthologues of these genes were not found, similar vertebrates genes found through BLAST (same PANTHER class) were used to infer the function  § GO function was missing and was inferred from other vertebrates orthologues | | | | |

**Table S2** Top 15 up- and down- regulated genes in the comparison of differential gene expressed between the larval and adult stage retinas of *P. amboinensis*.

**Table S3** Top 15 up- and down- regulated genes in the comparison of differential gene expressed between the juvenile and adult stage retinas of *P. amboinensis*.

|  | Gene ID | Gene name | logFC | Function |
| --- | --- | --- | --- | --- |
| Juvenile | CENPW | Centromere protein W | 4.750241851 | central nervous system development |
|  | KNL1 | Kinetochore scaffold 1 (Fragment) | 4.479773593 | cell division |
|  | esco2 | N-acetyltransferase ESCO2 | 4.378841758 | sister chromatid cohesion |
|  | smc2 | smc2 | 4.258458272 | mitotic chromosome condensation |
|  | LOC111584645 | Uncharacterized loci | 4.137472958 | NA |
|  | atoh7 | Protein atonal homolog 7 | 3.847819387 | retina development in camera-type eye |
|  | ldlra | Low density lipoprotein receptor | 3.845356476 | receptor-mediated endocytosis |
|  | birc5a | Baculoviral IAP repeat-containing 5a | 3.759776815 | cell cycle |
|  | anln | Anillin, actin-binding protein | 3.704461728 | mitotic nuclear division |
|  | e2f7 | Transcription factor E2F7 | 3.656453571 | transcription by RNA polymerase II |
|  | cks2 | Cyclin-dependent kinases regulatory subunit | 3.656375189 | regulation of mitotic cell cycle |
|  | tk1 | Thymidine kinase | 3.566563137 | DNA biosynthetic process |
|  | sstr2a* | Somatostatin receptor type 2 | 3.513830398 | neuropeptide signaling pathway |
|  | ncapg2 | Condensin-2 complex subunit G2 | 3.512886964 | cell division |
|  | fancd2 | FA complementation group D2 | 3.482178376 | cell cycle phase transition |
| Adult | LOC118469727 | Uncharacterized loci | -5.120580699 | NA |
|  | rxfp3 | Relaxin family peptide receptor 3 | -5.063394119 | G protein-coupled receptor signaling pathway |
|  | trim55b* | Tripartite motif-containing 55b | -4.719836882 | protein ubiquitination |
|  | gpr6* | G protein-coupled receptor 6 | -4.301728868 | G protein-coupled receptor signaling pathway |
|  | hpse2 | Heparanase 2 | -4.116951279 | extracellular matrix organization |
|  | zgc:154142 | Peptidase S1 domain-containing protein | -4.113645014 | proteolysis |
|  | rbp2a* | Cellular retinol-binding protein type II | -3.425592339 | vitamin A metabolic process |
|  | neflb | Neurofilament light polypeptide | -3.394532653 | intermediate filament bundle assembly |
|  | gck* | Phosphotransferase | -3.394197382 | glycolytic process |
|  | nppc | Natriuretic peptide C | -3.089320618 | cGMP biosynthetic process |
|  | opn1lw1* | Red-sensitive opsin-1 | -3.037198329 | visual perception |
|  | dcdc2b* | Doublecortin domain-containing 2B | -3.036608477 | intracellular signal transduction |
|  | rbp4l* | Purpurin | -2.88946946 | retinol transport |
|  | slc35a4* | Probable UDP-sugar transporter protein SLC35A4 | -2.814914809 | carbohydrate transport |
|  | zgc:101810 | Actin-related protein 2 | -2.801677398 | Arp2/3 complex-mediated actin nucleation |
| *these genes were not properly annotated and thus orthologues from zebrafish (Danio rerio) or medaka (Oryzias latipes) were used in Gene ID for GO Enrichement analysis  †orthologues of these genes were not found, similar vertebrates genes found through BLAST (same PANTHER class) were used to infer the function  § GO function was missing and was inferred from other vertebrates orthologues | | | | |

**Table S4** Top 15 up- and down- regulated genes in the comparison of differential gene expressed between the larval and juvenile stage retinas of *C. atripectoralis*.

|  | Gene ID | Gene name | logFC | Function |
| --- | --- | --- | --- | --- |
| Larvae | cryba2b | Crystallin, beta A2b | 19.41313252 | lens development in camera-type eye |
|  | cryba4 | Beta-crystallin A4 | 18.97828804 | lens development in camera-type eye |
|  | crygm1* | Crystallin, gamma M1 | 18.95800672 | lens development in camera-type eye |
|  | crygm3* | Crystallin, gamma M3 | 18.8252977 | lens development in camera-type eye |
|  | crygn2 | Gamma-crystallin N-B | 18.73273153 | lens development in camera-type eye |
|  | cryba1b* | Crystallin, beta A1b | 18.64184693 | lens development in camera-type eye |
|  | crygmxl2 | Crystallin, gamma MX, like 2 | 18.14560217 | lens development in camera-type eye |
|  | crygs3* | Crystallin, gamma S3 | 18.06528591 | lens development in camera-type eye |
|  | crybb1 | Beta-crystallin B1 | 17.09714102 | lens development in camera-type eye |
|  | crybb1l2 | Beta-crystallin B1 | 17.05134832 | lens development in camera-type eye |
|  | LOC111574450 | gamma-crystallin M3-like | 16.94823462 | lens development in camera-type eye |
|  | crygm2* | Crystallin, gamma M2 | 16.93976767 | lens development in camera-type eye |
|  | crygmxl1* | Crystallin, gamma MX, like 1 | 16.78458615 | lens development in camera-type eye |
|  | crygmx | crystallin, gamma MX | 16.70048157 | lens development in camera-type eye |
|  | crybb2 | Beta-crystallin B2 | 16.58954628 | lens development in camera-type eye |
| Juvenile | INHBC* | Inhibin beta B chain-like | -9.999303303 | SMAD protein signal transduction |
|  | opn1lw1* | Red-sensitive opsin-1 | -9.131991283 | visual perception |
|  | zmynd10 | Zinc finger, MYND-type containing 10 | -6.881446486 | positive regulation of motile cilium assembly |
|  | nfil3-5 | Nuclear factor, interleukin 3-regulated, member 5 | -6.641379763 | circadian rhythm |
|  | her11 | hairy-related 11 | -6.399424995 | regulation of neurogenesis |
|  | rorcb* | Nuclear receptor ROR-beta-like | -6.200450539 | regulation of transcription by RNA polymerase II |
|  | slc5a7a* | Solute carrier family 5 member 7a | -5.654734802 | regulation of neurotransmitter levels |
|  | mhc2b | Major histocompatibility complex class II integral membrane beta chain gene | -5.393520748 | immune response |
|  | hapln4 | hyaluronan and proteoglycan link protein 4 | -5.362782408 | central nervous system development |
|  | AVPI1 | Arginine vasopressin-induced protein 1 | -5.33567599 | cell cycle |
|  | NFIL3-2 | Nuclear factor, interleukin 3 regulated, member 2 | -5.250151826 | circadian rhythm |
|  | hrh3* | Histamine H3 receptor-like | -5.247799317 | chemical synaptic transmission |
|  | slc2a1* | Solute carrier family 2 (facilitated glucose transporter), member 1 | -5.150660723 | vitamin transport |
|  | LRAT† | Lecithin retinol acyltransferase-like | -5.071184143 | retinol metabolic process |
|  | pcdh2aa15* | Protocadherin beta-16 | -5.009596706 | cell adhesion |
| *these genes were not properly annotated and thus orthologues from zebrafish (Danio rerio) or medaka (Oryzias latipes) were used in Gene ID for GO Enrichement analysis  †orthologues of these genes were not found, similar vertebrates genes found through BLAST (same PANTHER class) were used to infer the function  § GO function was missing and was inferred from other vertebrates orthologues | | | | |

**Table S5** Top 15 up- and down- regulated genes in the comparison of differential gene expressed between the larval and adult stage retinas of *C. atripectoralis*.

|  | Gene ID | Gene name | logFC | Function |
| --- | --- | --- | --- | --- |
| Larvae | tnnt3b* | Troponin T type 3b | 25.80918067 | sarcomere organization |
|  | myhb* | myosin, heavy chain b | 25.45029187 | muscle contraction |
|  | pkmb | Pyruvate kinase | 23.82315285 | sprouting angiogenesis |
|  | cryba2b | Crystallin, beta A2b | 19.69199875 | lens development in camera-type eye |
|  | crygm1* | Crystallin, gamma M1 | 19.23687068 | lens development in camera-type eye |
|  | crygm3* | Crystallin, gamma M3 | 19.10416489 | lens development in camera-type eye |
|  | crygn2 | Gamma-crystallin N-B | 19.01159888 | lens development in camera-type eye |
|  | cryba1b* | Crystallin, beta A1b | 18.92071452 | lens development in camera-type eye |
|  | crybb1 | Beta-crystallin B1 | 18.91092848 | lens development in camera-type eye |
|  | crygmxl2 | Crystallin, gamma MX, like 2 | 18.42447051 | lens development in camera-type eye |
|  | crygs3* | Crystallin, gamma S3 | 18.34415403 | lens development in camera-type eye |
|  | crybb1l2 | Beta-crystallin B1 | 18.21525008 | lens development in camera-type eye |
|  | cryba4 | Beta-crystallin A4 | 17.39815555 | lens development in camera-type eye |
|  | LOC111574450 | gamma-crystallin M3-like | 17.22710458 | lens development in camera-type eye |
|  | crygm2* | Crystallin, gamma M2 | 17.21863427 | lens development in camera-type eye |
| Adults | INHBC* | Inhibin beta B chain-like | -10.22915908 | SMAD protein signal transduction |
|  | opn1lw1* | Red-sensitive opsin-1 | -9.37804641 | visual perception |
|  | nfil3-5 | Nuclear factor, interleukin 3-regulated, member 5 | -6.587254012 | circadian rhythm |
|  | hapln4 | hyaluronan and proteoglycan link protein 4 | -5.888774711 | central nervous system development |
|  | ccni2 | Cyclin I family, member 2 | -5.875206869 | mitotic cell cycle phase transition |
|  | rorcb* | Nuclear receptor ROR-beta-like | -5.8172308 | regulation of transcription by RNA polymerase II |
|  | NFIL3-2 | Nuclear factor, interleukin 3 regulated, member 2 | -5.689155624 | circadian rhythm |
|  | pcdh2aa15* | Protocadherin beta-16 | -5.643318115 | cell adhesion |
|  | cd74a§ | CD74 molecule, major histocompatibility complex, class II invariant chain a | -5.41189013 | positive regulation of cytokine-mediated signalling pathway |
|  | slc5a7a* | Solute carrier family 5 member 7a | -5.405548655 | regulation of neurotransmitter levels |
|  | mhc2b | Major histocompatibility complex class II integral membrane beta chain gene | -5.293254146 | immune response |
|  | frmd7* | FERM domain containing 7 | -5.245628238 | developmental process |
|  | slc5a5 | Solute carrier family 5 member 5 | -5.151152495 | sodium ion transport |
|  | zgc:172091* | GTPase IMAP family member 4-like | -5.105270575 | NA |
|  | her11 | hairy-related 11 | -5.061006486 | regulation of neurogenesis |
| *these genes were not properly annotated and thus orthologues from zebrafish (Danio rerio) or medaka (Oryzias latipes) were used in Gene ID for GO Enrichement analysis  †orthologues of these genes were not found, similar vertebrates genes found through BLAST (same PANTHER class) were used to infer the function  § GO function was missing and was inferred from other vertebrates orthologues | | | | |

**Table S6** Top 15 up- and down- regulated genes in the comparison of differential gene expressed between the juvenile and adult stage retinas of *C. atripectoralis*.

|  | Gene ID | Gene name | logFC | Function |
| --- | --- | --- | --- | --- |
| Juvenile | myhb* | myosin, heavy chain b | 25.80918067 | muscle contraction |
|  | tnnt3b* | Troponin T type 3b | 25.45029187 | sarcomere organization |
|  | pkmb | Pyruvate kinase | 23.82315285 | sprouting angiogenesis |
|  | myl1 | Myosin, light chain 1 | 19.69199875 | skeletal muscle tissue development |
|  | pvalb1* | Parvalbumin 1 | 19.23687068 | NA |
|  | ckma | Creatine kinase, muscle a | 19.10416489 | phosphorylation |
|  | acta1b† | Actin alpha 1, skeletal muscle b | 19.01159888 | skeletal muscle fiber development |
|  | clic5b | Chloride intracellular channel 5b | 18.92071452 | monoatomic ion transport |
|  | ckmb† | Creatine kinase, muscle b | 18.91092848 | phosphorylation |
|  | desma | Desmin a | 18.42447051 | skeletal muscle organ development |
|  | pvalb3§ | Parvalbumin 3 | 18.34415403 | excitatory chemical synaptic transmission |
|  | myhz1.1* | Myosin, heavy polypeptide 1.1 | 18.21525008 | muscle contraction |
|  | atp2a1* | ATPase sarcoplasmic/endoplasmic reticulum Ca2+ transporting 1 | 17.39815555 | calcium ion transmembrane transport |
|  | mafk | V-maf avian musculoaponeurotic fibrosarcoma oncogene homolog K | 17.22710458 | regulation of transcription by RNA polymerase II |
|  | zgc:153704 | Lipocalin-like | 17.21863427 | NA |
| Adults | epor | Erythropoietin receptor | -10.22915908 | cytokine-mediated signaling pathway |
|  | tmod1 | tropomodulin 1 | -9.37804641 | muscle contraction |
|  | wnk1a* | WNK lysine deficient protein kinase 1a | -6.587254012 | angiogenesis |
|  | kcnmb4* | Potassium calcium-activated channel subfamily M regulatory beta subunit 4 | -5.888774711 | detection of stimulus |
|  | synpo2* | synaptopodin 2 | -5.875206869 | skeletal muscle tissue development |
|  | trim46a | tripartite motif containing 46a | -5.8172308 | neuron migration |
|  | si:dkey-219c3.2* | TAFH domain-containing protein | -5.689155624 | DNA-templated transcription, initiation |
|  | zgc:171704§ | Ras-related and estrogen-regulated growth inhibitor-like protein | -5.643318115 | positive regulation of transcription by RNA polymerase I |
|  | ttc39b† | Tetratricopeptide repeat domain 39B | -5.41189013 | regulation of transport |
|  | ppfibp1a* | PPFIA binding protein 1a | -5.405548655 | neuromuscular junction development |
|  | znf644b | Zinc finger protein 644b | -5.293254146 | regulation of transcription by RNA polymerase II |
|  | galnt9* | Polypeptide N-acetylgalactosaminyltransferase 9 | -5.245628238 | protein O-linked glycosylation |
|  | whrnb§ | Whirlin b | -5.151152495 | retina homeostasis |
|  | plekha1b§ | Pleckstrin homology domain containing, family A member 1b | -5.105270575 | NA |
|  | si:ch211-276i12.4 | uncharacterized si:ch211-276i12.4 | -5.061006486 | NA |
| *these genes were not properly annotated and thus orthologues from zebrafish (Danio rerio) or medaka (Oryzias latipes) were used in Gene ID for GO Enrichement analysis  †orthologues of these genes were not found, similar vertebrates genes found through BLAST (same PANTHER class) were used to infer the function  § GO function was missing and was inferred from other vertebrates orthologues | | | | |

**Table S7** Top 15 up- and down- regulated genes in the comparison of differential gene expressed between the larvae and juvenile stage retinas of *A. akindynos*

|  | Gene ID | Gene name | logFC | Function |
| --- | --- | --- | --- | --- |
| Larvae | crybb1 | Crystallin, beta B1 | 19.53054 | lens development in camera-type eye |
|  | cryba1b* | Crystallin, beta A1b | 18.87027856 | lens development in camera-type eye |
|  | crygm1* | Crystallin, gamma M1 | 18.81490769 | lens development in camera-type eye |
|  | cryba4 | Crystallin, beta A4 | 18.78383673 | lens development in camera-type eye |
|  | crygm3* | Crystallin, gamma M3 | 18.6773972 | lens development in camera-type eye |
|  | LOC111574699 | Gamma-crystallin M3-like | 18.44011453 | lens development in camera-type eye |
|  | crygmxl2 | Crystallin, gamma MX, like 2 | 18.33564485 | lens development in camera-type eye |
|  | crygs3* | Crystallin, gamma S3 | 18.28162284 | lens development in camera-type eye |
|  | LOC111586589 | Gamma-crystallin M3-like | 18.21756424 | lens development in camera-type eye |
|  | LOC111586605 | Gamma-crystallin M3-like | 18.07742692 | lens development in camera-type eye |
|  | LOC111586595 | Gamma-crystallin M3-like | 17.85294402 | lens development in camera-type eye |
|  | crybgx | Crystallin beta gamma X | 17.43467265 | lens development in camera-type eye |
|  | LOC111574450 | Gamma-crystallin M3-like | 17.21651781 | lens development in camera-type eye |
|  | crygmx | crystallin, gamma MX | 16.9797586 | lens development in camera-type eye |
|  | LOC111586599 | Gamma-crystallin M3-like | 16.88214862 | lens development in camera-type eye |
| Juvenile | opn1sw2 | Opsin 1 (cone pigments), short-wave-sensitive 2 | -8.20266873 | visual perception |
|  | her11 | Hairy-related 11 | -7.49246285 | regulation of neurogenesis |
|  | INHBC* | Inhibin beta B chain-like | -7.334487499 | SMAD protein signal transduction |
|  | LOC118470026 | uncharacterized loci | -7.016138179 | NA |
|  | rad9b | RAD9 checkpoint clamp component B | -6.980511854 | DNA repair |
|  | LOC118471372 | uncharacterized loci | -6.963514616 | NA |
|  | LOC129348562 | uncharacterized loci | -6.775357088 | NA |
|  | fbxw10§ | F-box and WD repeat domain containing 10 | -6.33254029 | G1/S transition of mitotic cell cycle |
|  | nfil3-5 | Nuclear factor, interleukin 3-regulated, member 5 | -6.188526536 | circadian rhythm |
|  | rorcb* | Nuclear receptor ROR-beta-like | -6.152761516 | regulation of transcription by RNA polymerase II |
|  | GUCY2CA | Guanylate cyclase 2Ca | -6.082281369 | cGMP biosynthetic process |
|  | LOC129350040 | uncharacterized loci | -6.022913226 | NA |
|  | slc19a3a | Solute carrier family 19 member 3a | -5.951366991 | vitamin transport |
|  | a2ml* | Alpha-2-macroglobulin-like | -5.693515473 | NA |
|  | LOC118471463 | uncharacterized loci | -5.625392146 | NA |
| *these genes were not properly annotated and thus orthologues from zebrafish (Danio rerio) or medaka (Oryzias latipes) were used in Gene ID for GO Enrichement analysis  †orthologues of these genes were not found, similar vertebrates genes found through BLAST (same PANTHER class) were used to infer the function  § GO function was missing and was inferred from other vertebrates orthologues | | | | |

**Table S8** Top 15 up- and down- regulated genes in the comparison of differential gene expressed between the larvae and adults stage retinas of *A. akindynos*

|  | Gene ID | Gene name | logFC | Function |
| --- | --- | --- | --- | --- |
| Larvae | crybb1 | Crystallin, beta B1 | 19.49371037 | lens development in camera-type eye |
|  | cryba2b | Crystallin, beta A2b | 19.13917838 | lens development in camera-type eye |
|  | cryba1b* | Crystallin, beta A1b | 18.83346991 | lens development in camera-type eye |
|  | crygm1* | Crystallin, gamma M1 | 18.77812453 | lens development in camera-type eye |
|  | crygm3* | Crystallin, gamma M3 | 18.64063618 | lens development in camera-type eye |
|  | LOC111574699 | Gamma-crystallin M3-like | 18.40330882 | lens development in camera-type eye |
|  | crygmxl2 | Crystallin, gamma MX, like 2 | 18.29889801 | lens development in camera-type eye |
|  | LOC111574368 | Gamma-crystallin M3-like | 18.24482117 | lens development in camera-type eye |
|  | LOC111586589 | Gamma-crystallin M3-like | 18.18079353 | lens development in camera-type eye |
|  | cryba1l1 | Crystallin, beta A1, like 1 | 18.10016483 | lens development in camera-type eye |
|  | LOC111586605 | Gamma-crystallin M3-like | 18.04064127 | lens development in camera-type eye |
|  | LOC111586595 | Gamma-crystallin M3-like | 17.81615711 | lens development in camera-type eye |
|  | cryba4 | Crystallin, beta A4 | 17.78526825 | lens development in camera-type eye |
|  | LOC111574450 | Gamma-crystallin M3-like | 17.1797724 | lens development in camera-type eye |
|  | LOC111575220 | Gamma-crystallin M2-like | 17.06851945 | lens development in camera-type eye |
| Adult | opn1sw2 | Opsin 1 (cone pigments), short-wave-sensitive 2 | -8.799370959 | visual perception |
|  | rad9b | RAD9 checkpoint clamp component B | -7.653378028 | DNA repair |
|  | INHBC† | Inhibin beta B chain-like | -7.631971919 | signal transduction |
|  | LOC118470026 | uncharacterized | -7.030708942 | NA |
|  | fbxw10§ | F-box and WD repeat domain containing 10 | -6.725011099 | G1/S transition of mitotic cell cycle |
|  | LOC118471372 | uncharacterized | -6.503777655 | NA |
|  | MUC19* | Mucin 19 | -6.44990364 | hematopoietic progenitor cell differentiation |
|  | ighm† | Immunoglobulin heavy constant mu | -6.086969247 | MAPK cascade |
|  | SI:CH1073-416J23.1 | Rac GTPase-activating protein 1 | -6.052300653 | animal organ development |
|  | nfil3-5 | Nuclear factor, interleukin 3-regulated, member 5 | -5.862724551 | circadian rhythm |
|  | LOC118471091 | uncharacterized | -5.805312108 | NA |
|  | LOC111571222 | uncharacterized | -5.790167356 | NA |
|  | LOC129347667† | Immunoglobulin mu heavy chain-like | -5.759730315 | immune response |
|  | LOC111577565† | Ig kappa chain V region Mem5-like | -5.742063915 | immune response |
|  | otol1b | Otolin 1b | -5.703894506 | extracellular matrix organization |
| *these genes were not properly annotated and thus orthologues from zebrafish (Danio rerio) or medaka (Oryzias latipes) were used in Gene ID for GO Enrichement analysis  †orthologues of these genes were not found, similar vertebrates genes found through BLAST (same PANTHER class) were used to infer the function  § GO function was missing and was inferred from other vertebrates orthologues | | | | |

**Table S9** Top 15 up- and down- regulated genes in the comparison of differential gene expressed between the juvenile and adult stage retinas of *A. akindynos*

|  | Gene ID | Gene name | logFC | Function |
| --- | --- | --- | --- | --- |
| Juvenile | gys2 | Glycogen synthase 2 | 5.06164804 | glycogen biosynthetic process |
|  | rapsn | Receptor-associated protein of the synapse, 43kD | 4.968618006 | acetylcholine receptor binding |
|  | acanb | Aggrecan b | 4.61878326 | central nervous system development |
|  | si:dkey-219e21.4* | Tripartite motif-containing protein 14-like | 4.401272326 | protein ubiquitination |
|  | si:ch1073-324l1.1* | Tripartite motif-containing protein 35-like | 4.301096151 | protein ubiquitination |
|  | enkur* | Enkurin, TRPC channel interacting protein | 4.21533214 | NA |
|  | daam1a* | Dishevelled associated activator of morphogenesis 1a | 4.135068245 | actin cytoskeleton organizatio |
|  | tmem45b§ | Transmembrane protein 45B | 4.02817397 | immune system process |
|  | hykk | Hydroxylysine kinase | 3.821109274 | NA |
|  | bora | Protein aurora borealis | 3.820283538 | cell cycle |
|  | GJB8 | Gap junction protein beta 8 | 3.817563161 | cell communication |
|  | uba52* | Ubiquitin A-52 residue ribosomal protein fusion product 1 | 3.444537974 | protein ubiquitination |
|  | sult1st3* | Sulfotransferase family 1, cytosolic sulfotransferase 3 | 3.351959282 | NA |
|  | igf2bp1 | Insulin-like growth factor 2 mRNA binding protein 1 | 3.351375954 | neural retina development |
|  | desma | Desmin a | 3.288335708 | skeletal muscle organ development |
| Adult | scpp7 | Secretory calcium-binding phosphoprotein 7 | -7.157005181 | NA |
|  | txnipb* | Thioredoxin-interacting protein | -6.879087519 | protein transport |
|  | mpl | MPL proto-oncogene, thrombopoietin receptor | -6.740409392 | cytokine-mediated signaling pathway |
|  | ETSRP | ETS1-related protein | -6.566660977 | angiogenesis |
|  | foxf2a | Forkhead box F2a | -6.56479969 | animal organ morphogenesis |
|  | ano1a* | Anoctamin 1, calcium activated chloride channel a | -6.511448475 | chloride transmembrane transport |
|  | tbx18 | T-box transcription factor 18 | -6.42935784 | cell fate specification |
|  | SI:CH211-286O17.1 | Hematopoietic progenitor cell antigen CD34 | -6.243966852 | NA |
|  | Cldn4† | Claudin 4 | -6.22764606 | Ion transport |
|  | mmp13a | Matrix metallopeptidase 13a | -6.118044837 | catabolic process |
|  | gask1b | Golgi associated kinase 1B | -6.004585105 | NA |
|  | capn2l* | Calcium-activated neutral proteinase 2 | -5.939366602 | proteolysis |
|  | hdr | Hematopoietic death receptor | -5.889518199 | positive regulation of apoptotic process |
|  | col8a1b* | Collagen, type VIII, alpha 1b | -5.863118185 | extracellular matrix organization |
|  | SI:CH73-52P7.1 | uncharacterized | -5.848586246 | NA |
| *these genes were not properly annotated and thus orthologues from zebrafish (Danio rerio) or medaka (Oryzias latipes) were used in Gene ID for GO Enrichement analysis  †orthologues of these genes were not found, similar vertebrates genes found through BLAST (same PANTHER class) were used to infer the function  § GO function was missing and was inferred from other vertebrates orthologues | | | | |

**Table S10** Presence (denoted by a X) of complex UV patterns and simple UV patterns in three ontogenetic stages of the eleven species of damselfish investigated.

| Species | Stage | Complex UV patterns | Simple UV Patterns |
| --- | --- | --- | --- |
| *Chromis atripectoralis* | Larval | NA | NA |
|  | Juvenile |  | X |
|  | Adult |  | X |
| *Dascyllus aruanus* | Larval |  |  |
|  | Juvenile |  | X |
|  | Adult |  | X |
| *Dischistodus perspicillatus* | Larval |  |  |
|  | Juvenile |  |  |
|  | Sub-adults | X |  |
|  | Adult | X |  |
| *Chrysiptera flavipinnis* | Larval | NA | NA |
|  | Juvenile |  | X |
|  | Adult |  | X |
| *Neoglyphidodon melas* | Larval | NA | NA |
|  | Juvenile |  | X |
|  | Adult |  |  |
| *Amphiprion akindynos* | Larval |  |  |
|  | Juvenile |  | X |
|  | Adult |  | X |
| *Pomacentrus pavo* | Adult | NA | NA |
|  | Juvenile | X |  |
|  | Adult | X |  |
| *Pomacentrus bankanensis* | Larval |  |  |
|  | Juvenile | X |  |
|  | Adult | X |  |
| *Pomacentrus chrysurus* | Larval |  |  |
|  | Juvenile | X |  |
|  | Adult |  |  |
| *Pomacentrus amboinensis* | Larval |  |  |
|  | Juvenile | X |  |
|  | Adult | X |  |
| *Pomacentrus nagasakiensis* | Larval | NA | NA |
|  | Juvenile | X |  |
|  | Adult | X |  |

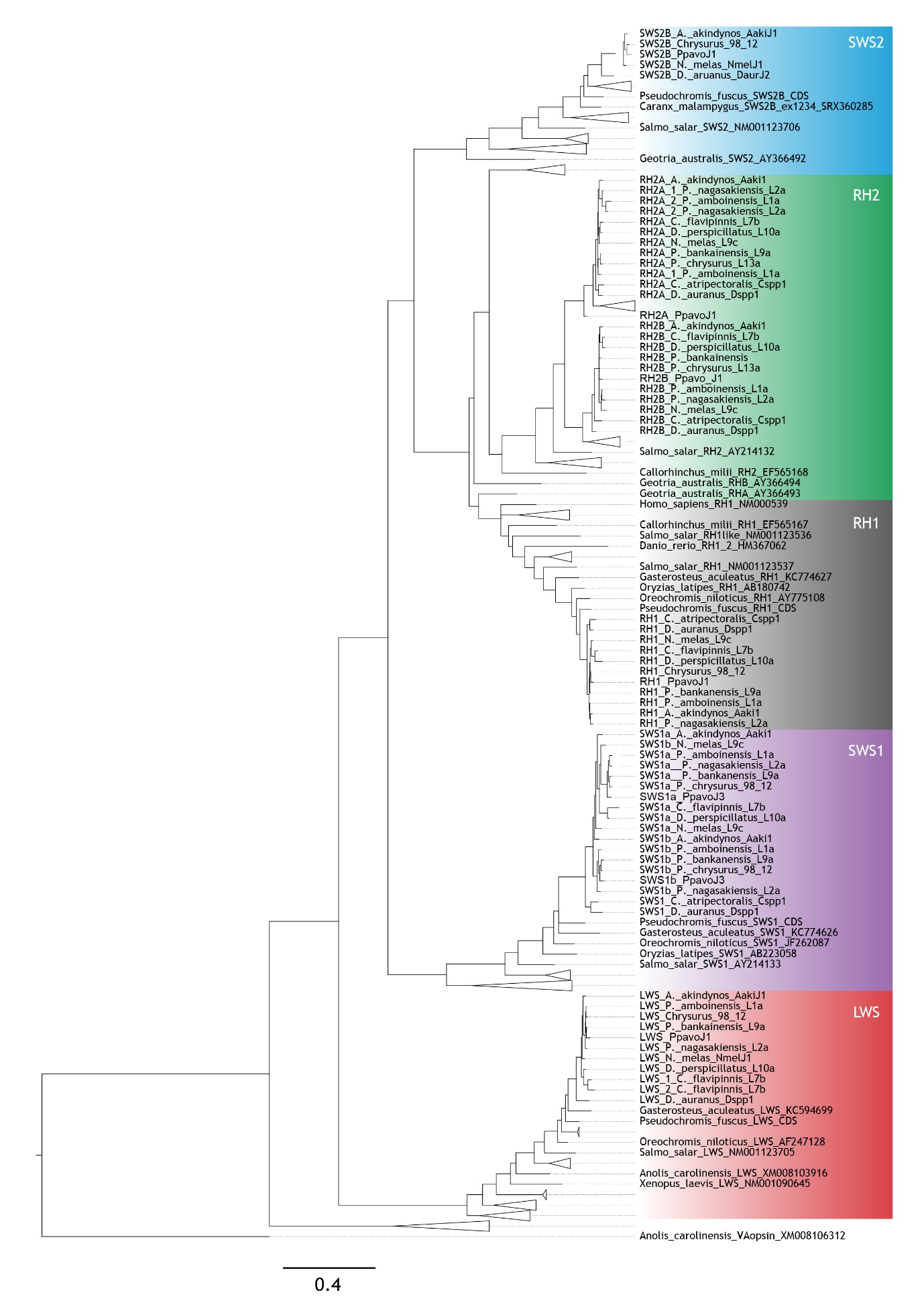

**Figure S1** Phylogenetic reconstruction of all opsins observed in the transcriptomes of the eleven species of damselfish family tageted in this study.

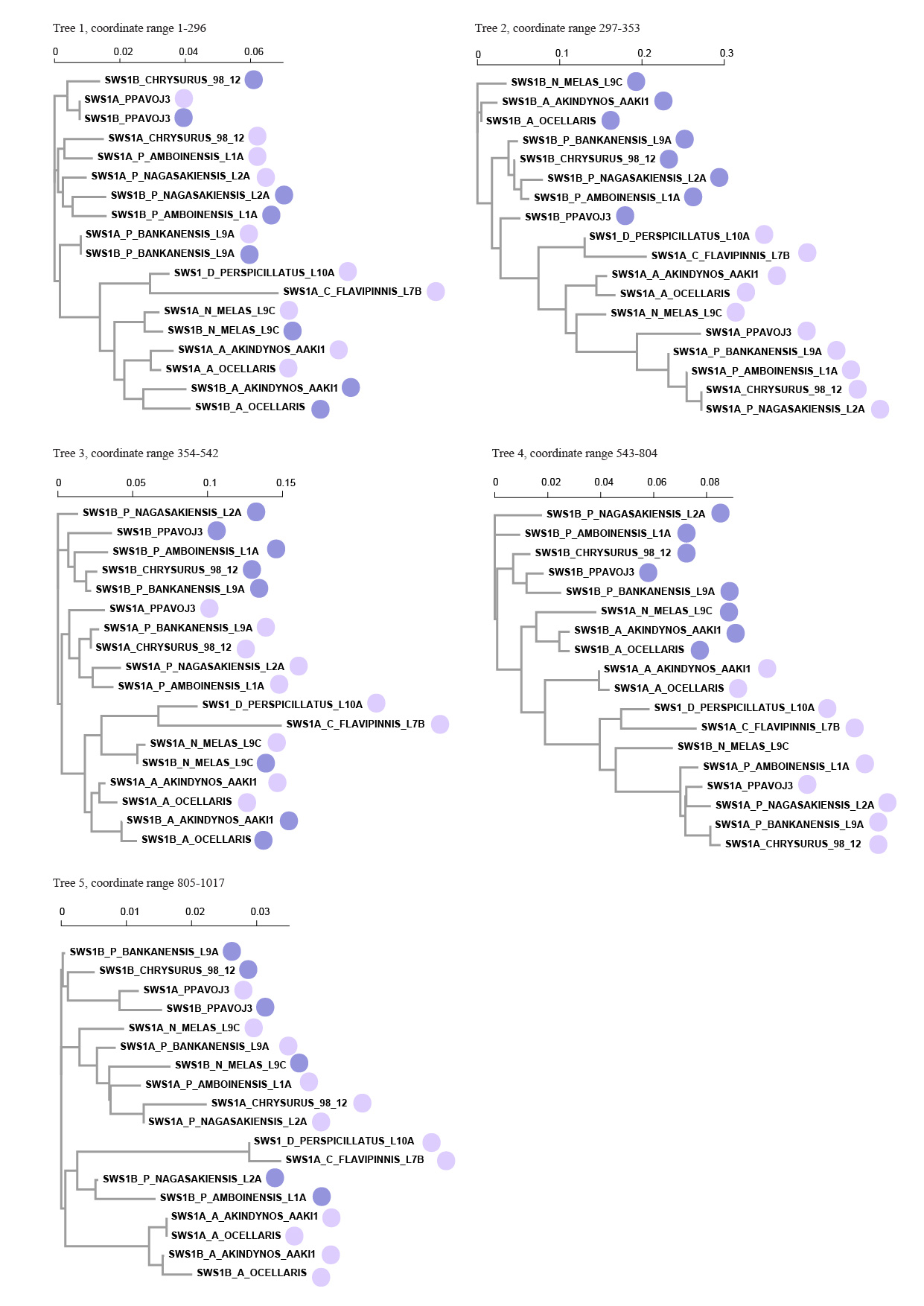

**Figure S2** GARD analysis of the *sws1* orthologues and paralogues in the species expressing *sws1α* and *sws1β*.

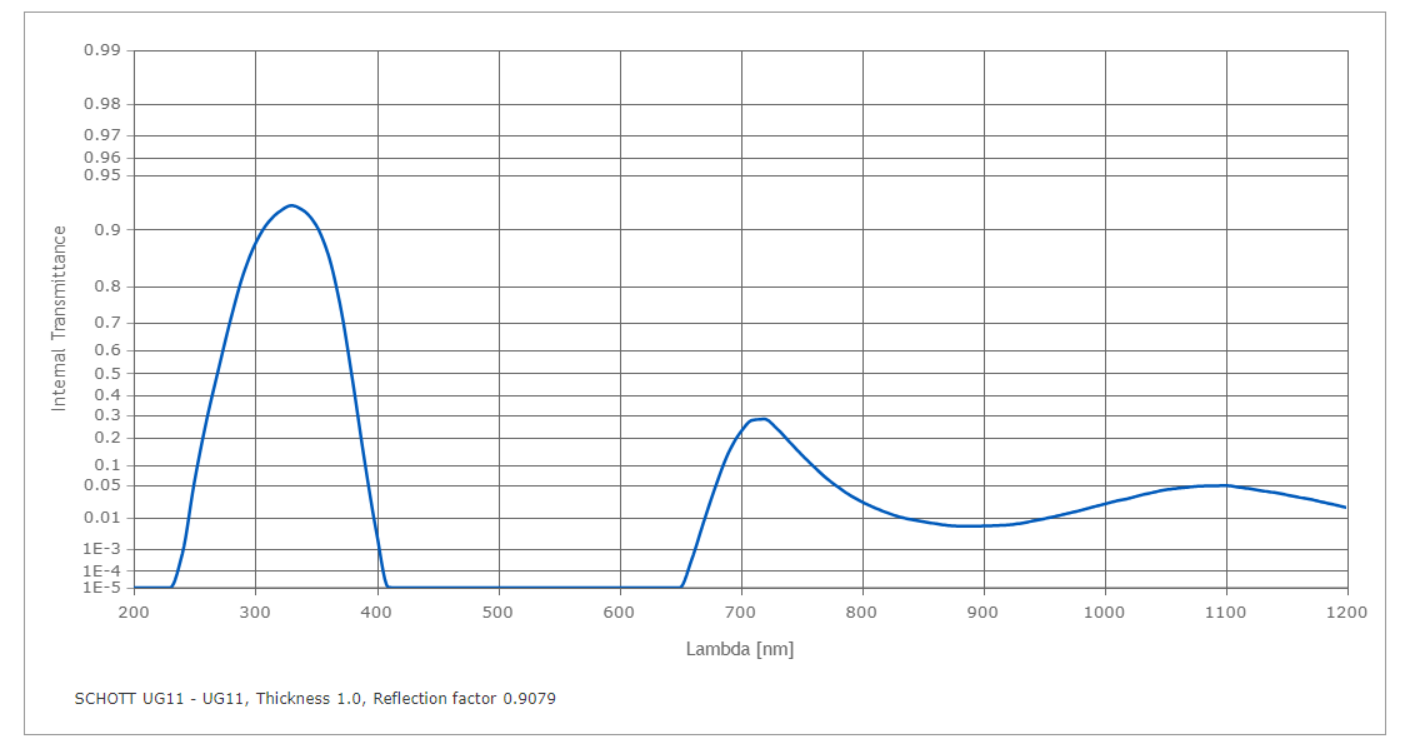

**Figure S3** Transmission of the ultraviolet (UV) bandpass filter Schott UG11 visible-blocking.

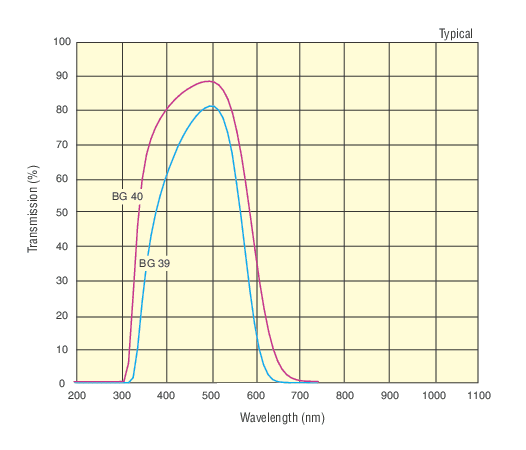

**Figure S4** Transmittance of Newport FSQ-BG39 blue bandpass filter.

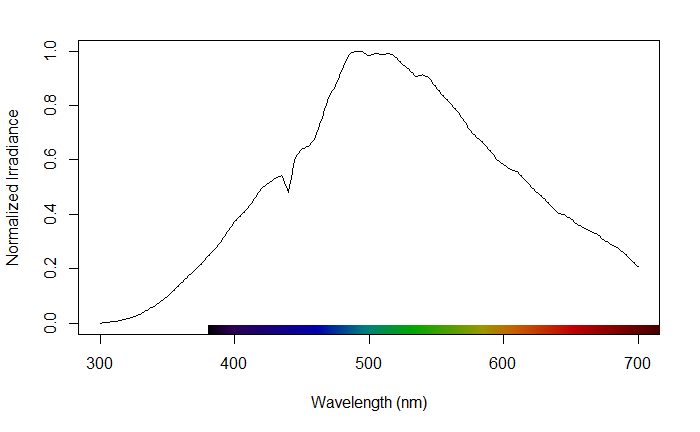

**Figure S5** Spectrum of the Nikon Speedlight SB-26 flashes.

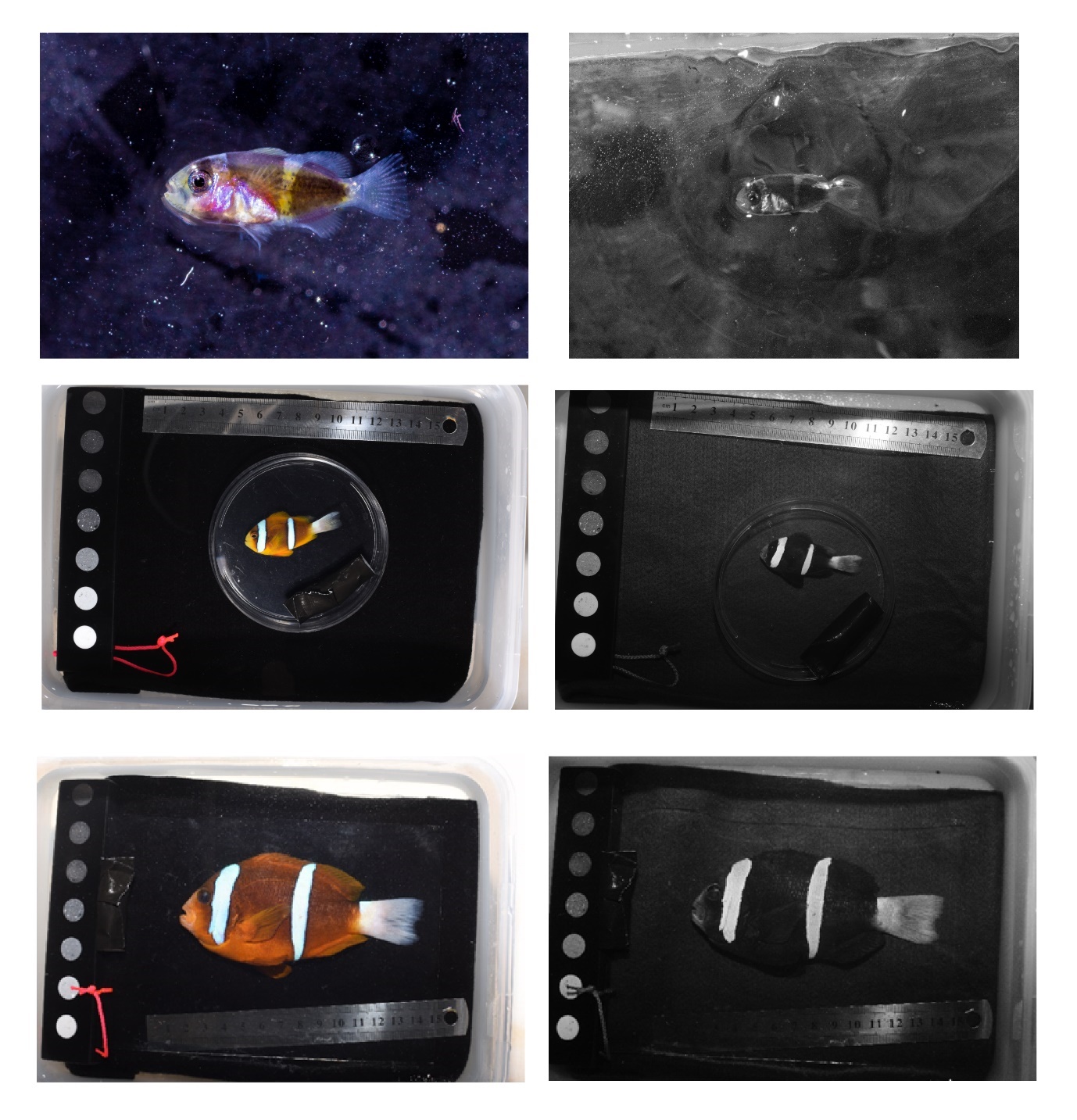

**Figure S6** Amphiprion akindynos pictures taken both with a stock (left) and an ultraviolet camera (right). From the top: larval, juvenile and adult stages.

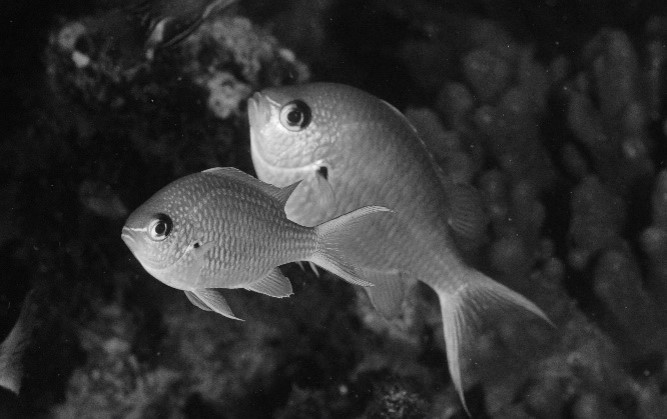

**Figure S7** Juvenile and adult Chromis atripectoralis. Image taken with an underwater ultraviolet camera setup.

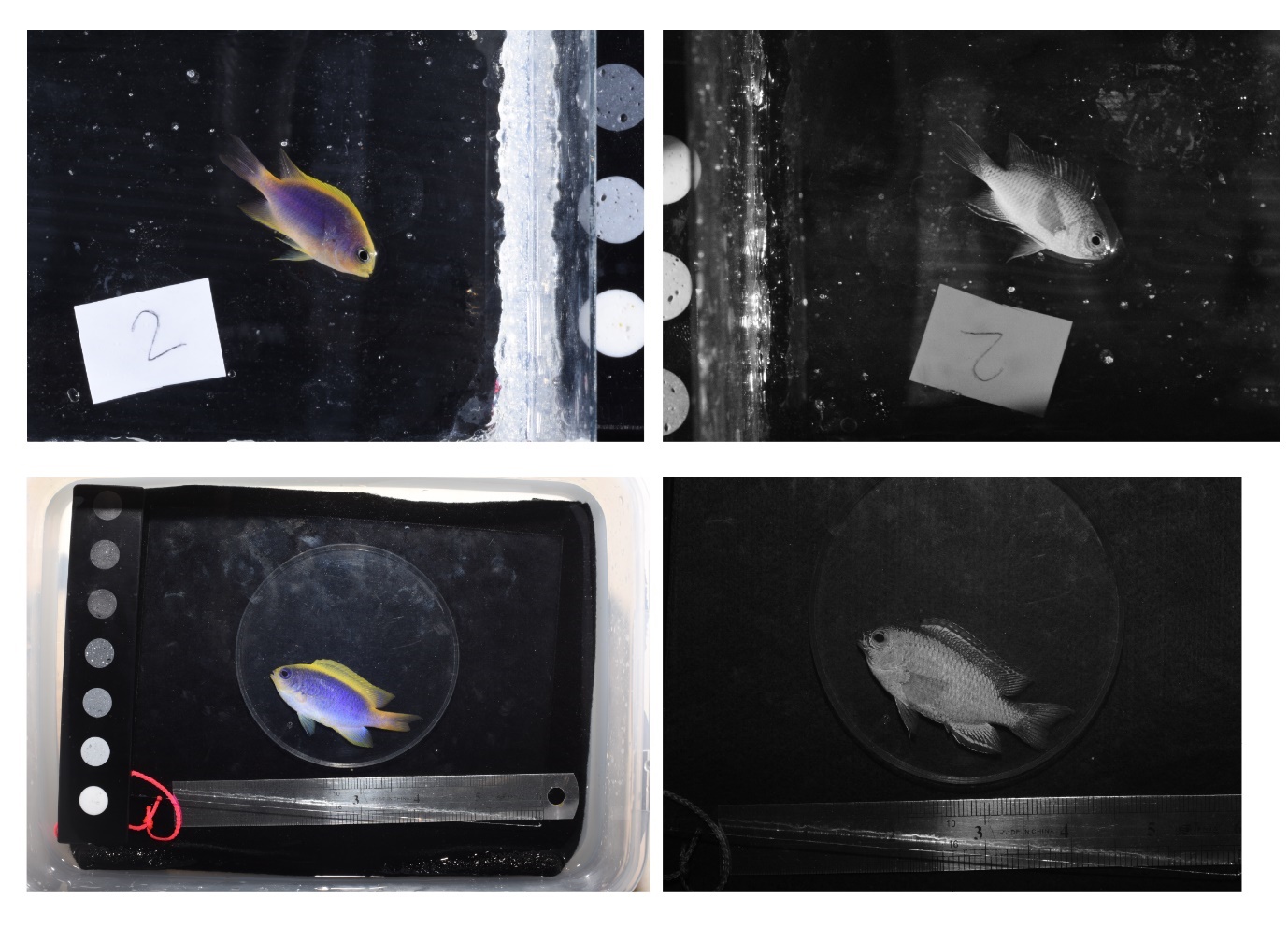

**Figure S8** Chrysiptera flavipinnis pictures taken both with a stock (left) and an ultraviolet camera (right). From the top: juvenile and adult stages.

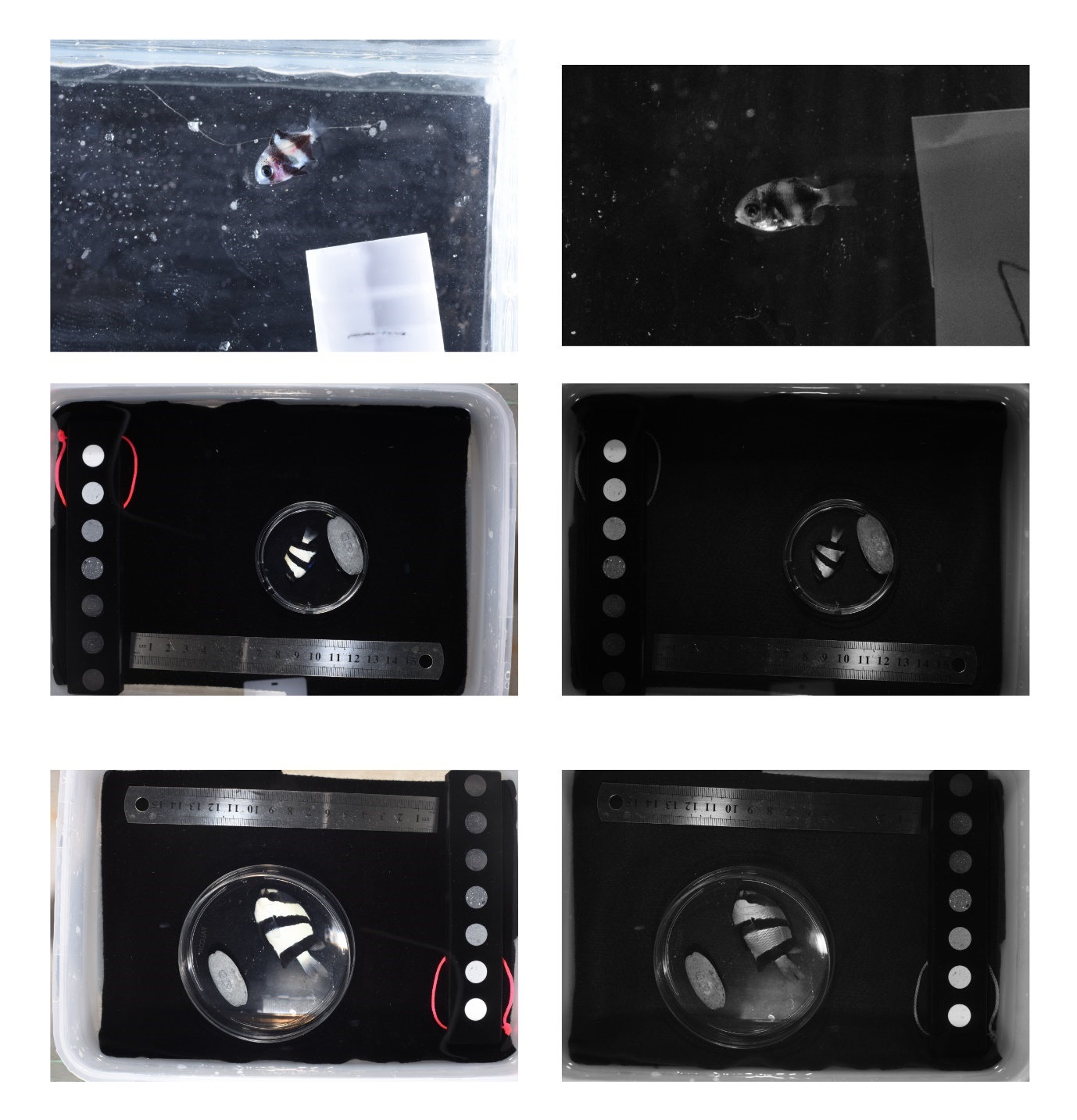

**Figure S9** Dascyllus aruanus pictures taken both with a stock (left) and an ultraviolet camera (right). From the top: larval, juvenile and adult stages.

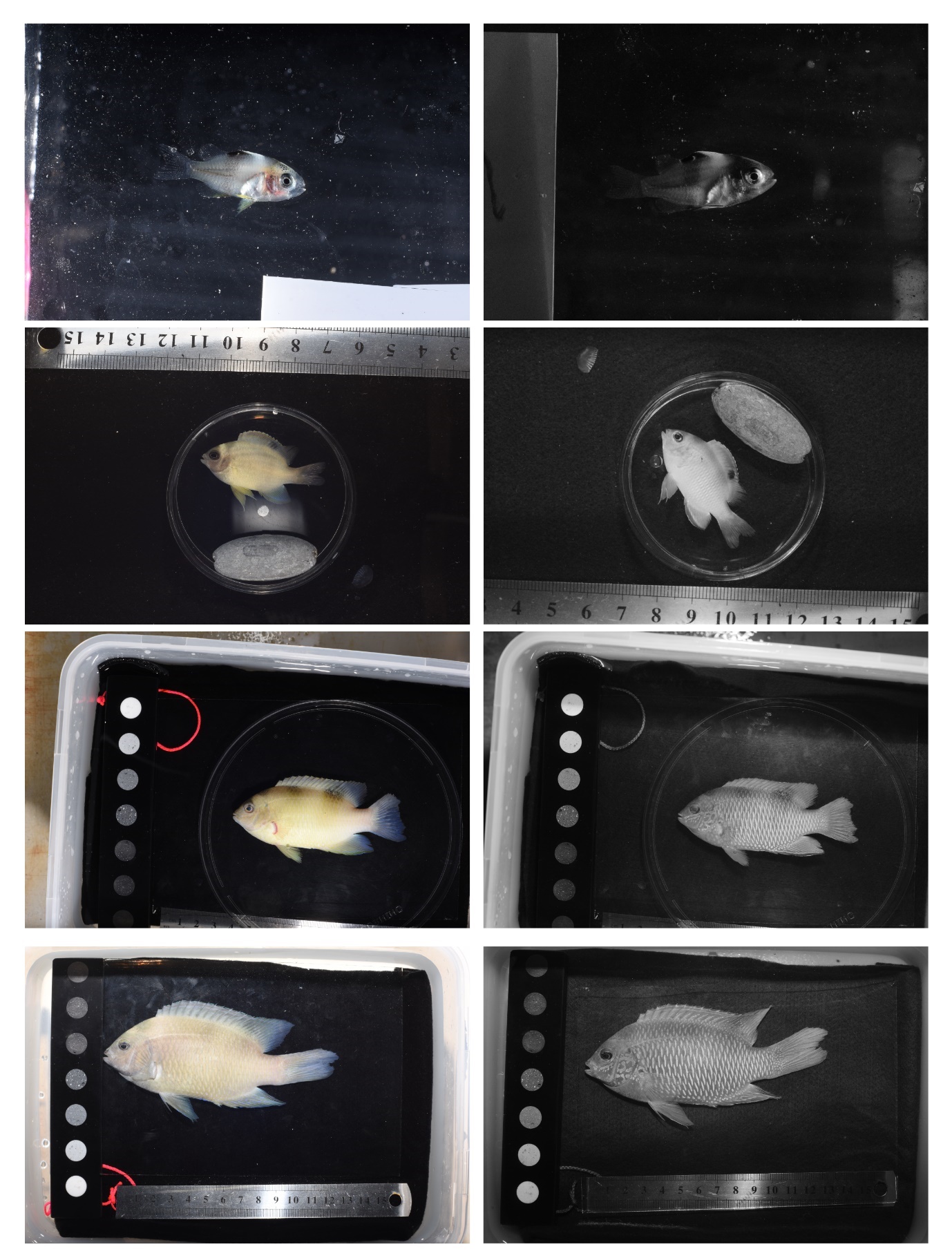

**Figure S10** Dischistodus perspicillatus pictures taken both with a stock (left) and an ultraviolet camera (right). From the top: larval, juvenile, sub-adult, and adult stages.

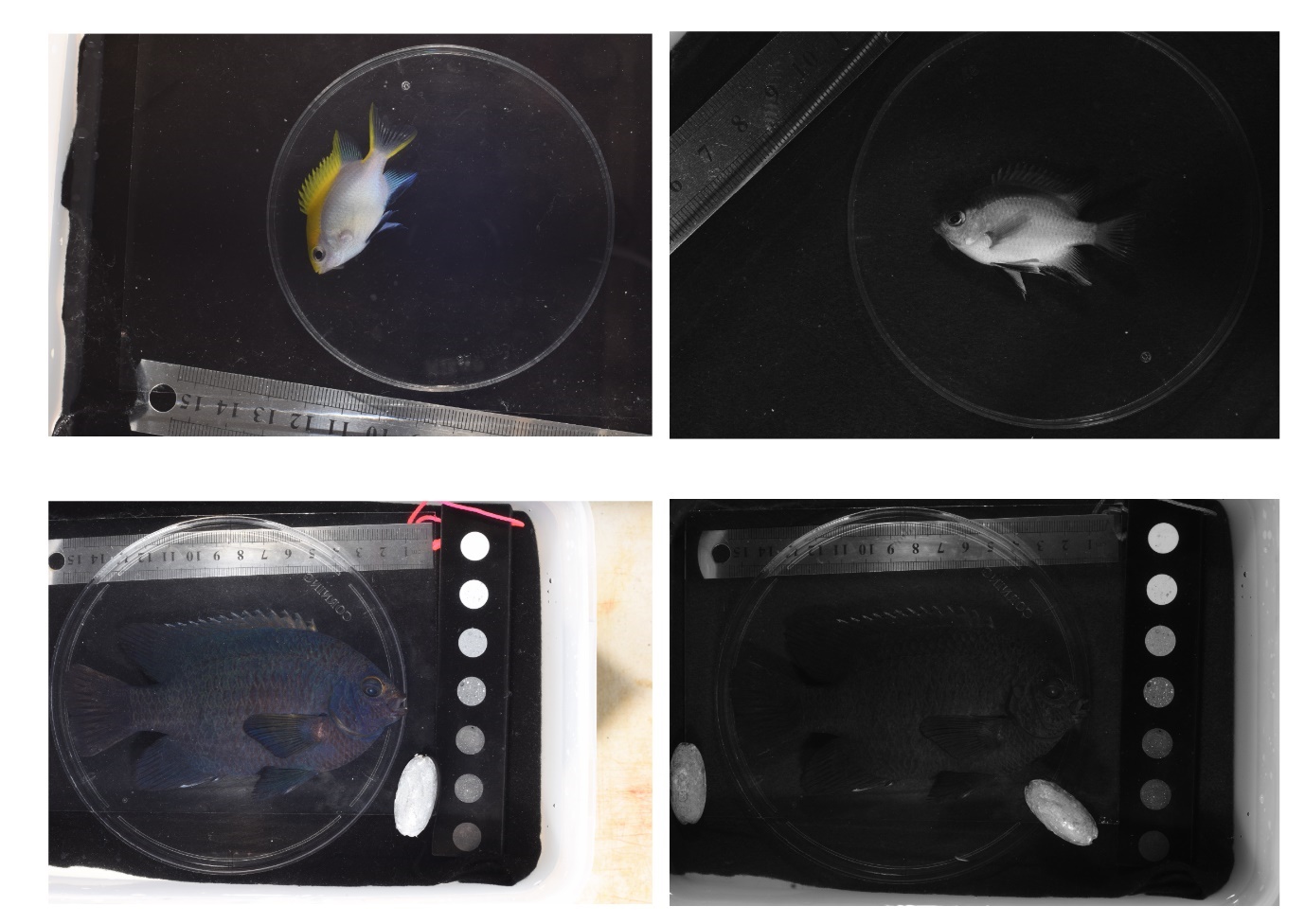

**Figure S11** Neoglyphidodon melas pictures taken both with a stock (left) and an ultraviolet camera (right). From the top: juvenile, and adult stages.

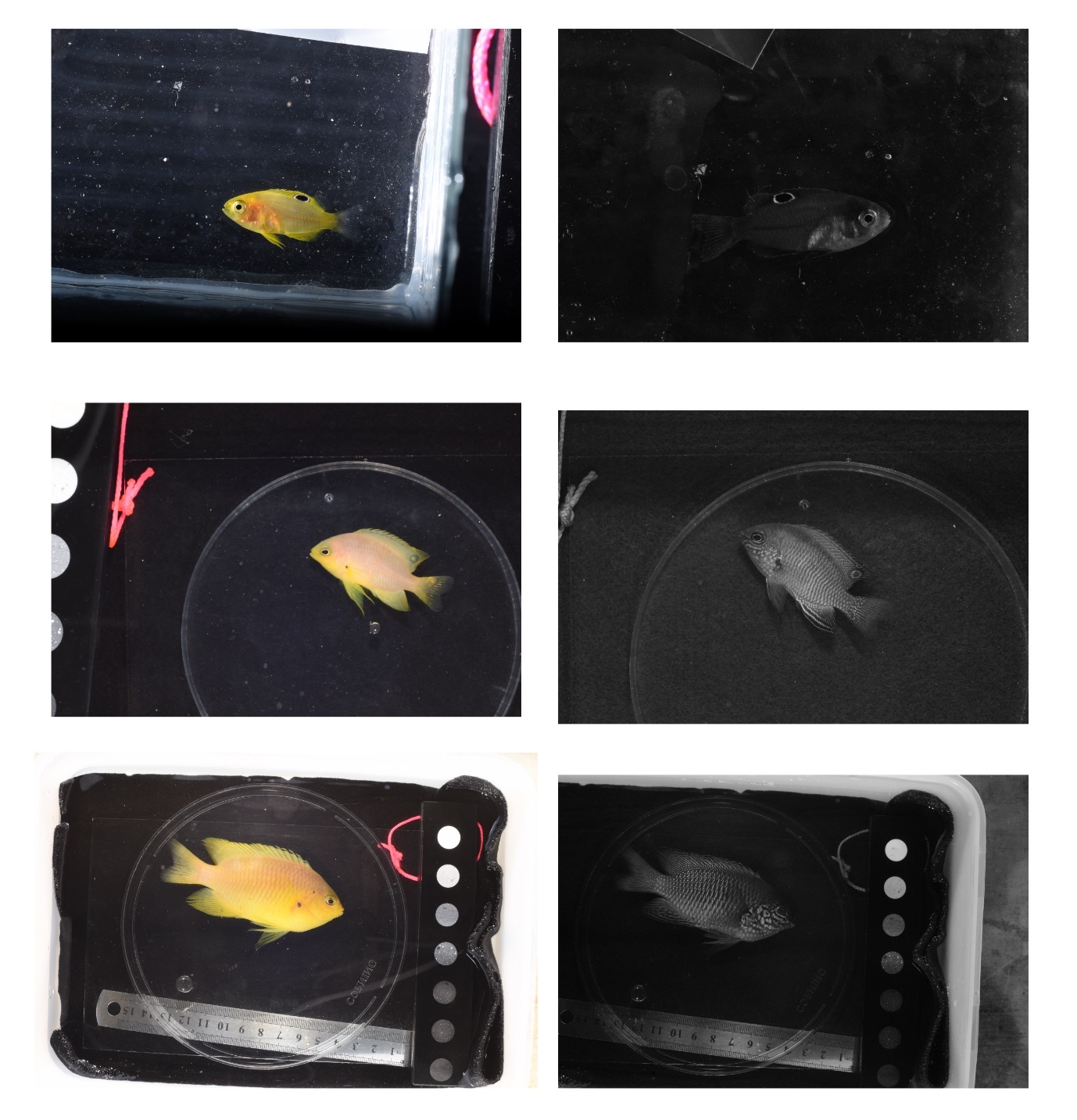

**Figure S12** Pomacentrus amboinensis pictures taken both with a stock (left) and an ultraviolet camera (right). From the top: larval, juvenile, and adult stages.

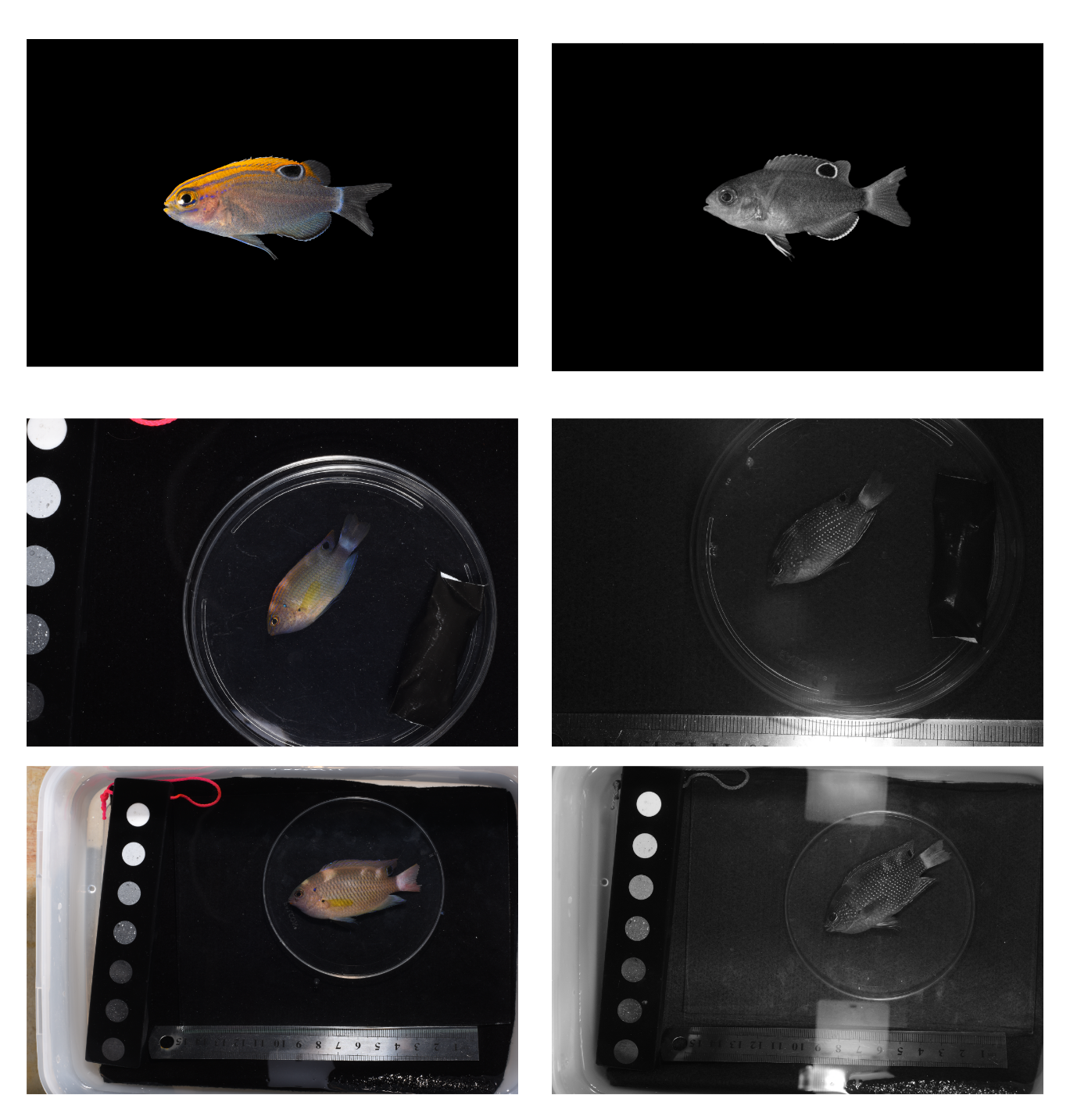

**Figure S13** Pomacentrus bankanensis pictures taken both with a stock (left) and an ultraviolet camera (right). From the top: larval, juvenile, and adult stages.

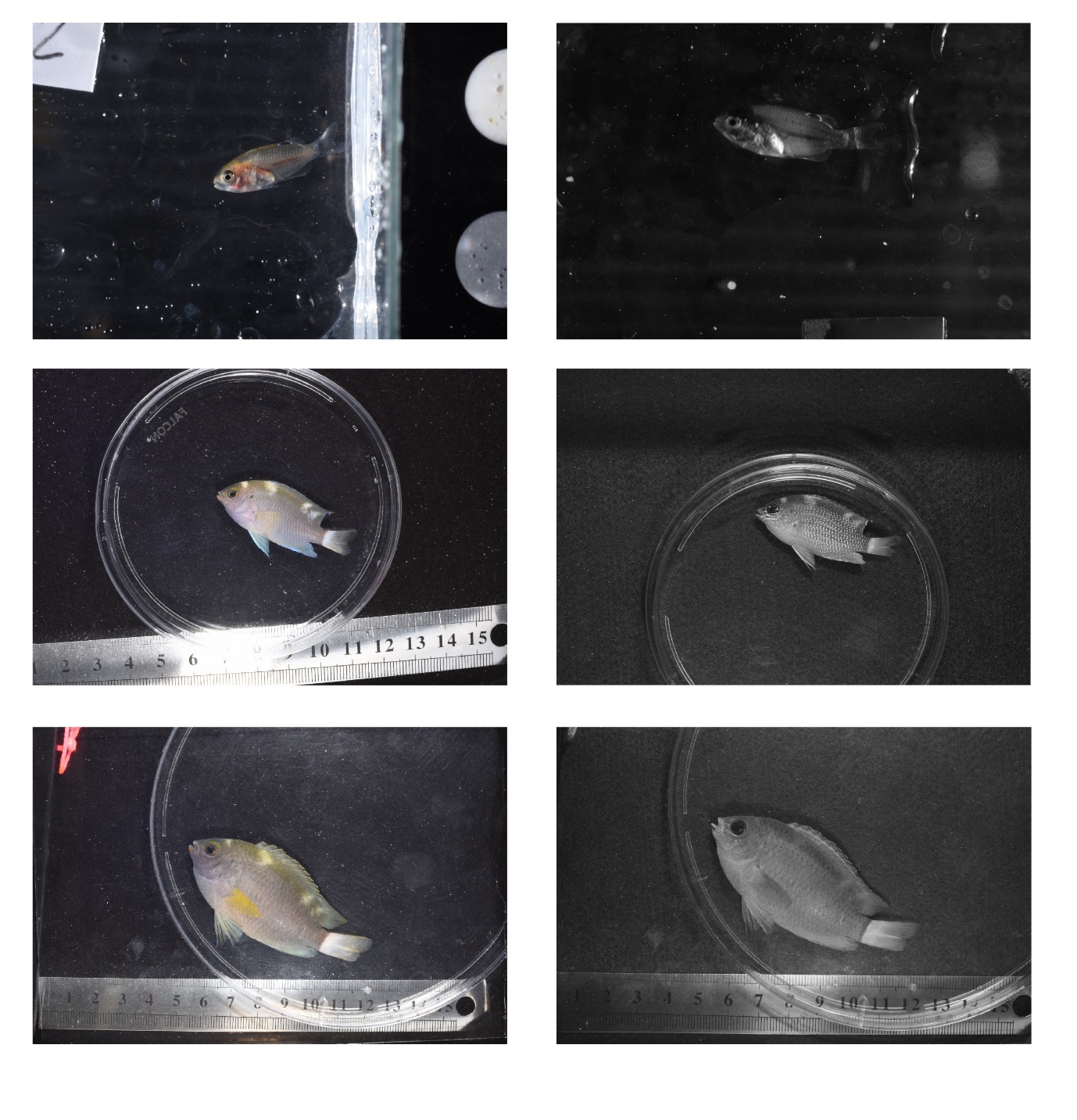

**Figure S14** Pomacentrus chrysurus pictures taken both with a stock (left) and an ultraviolet camera (right). From the top: larval, juvenile, and adult stages.

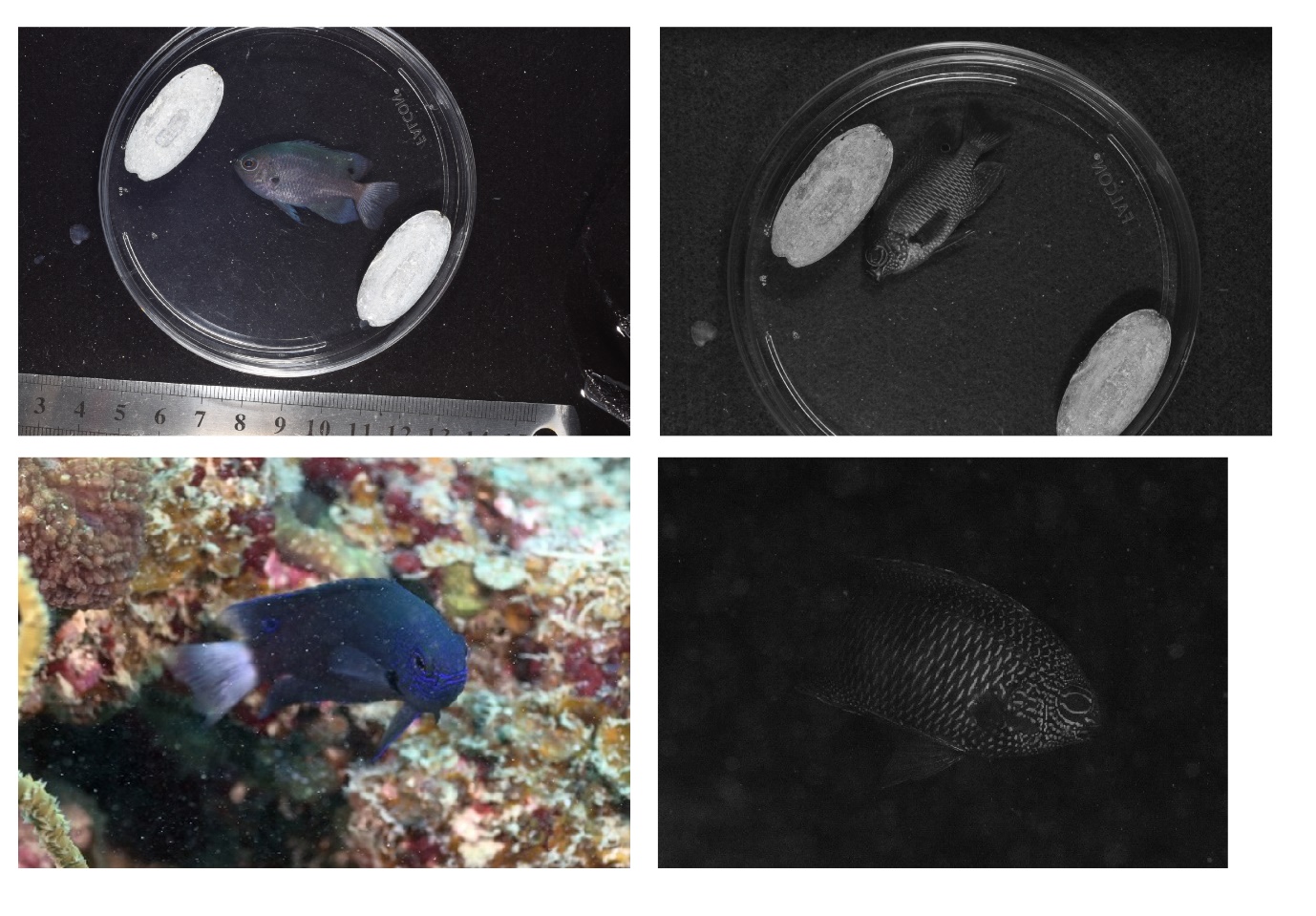

**Figure S15** Pomacentrus nagasakiensis pictures taken both with a stock (left) and an ultraviolet camera (right). From the top: juvenile, and adult stages.

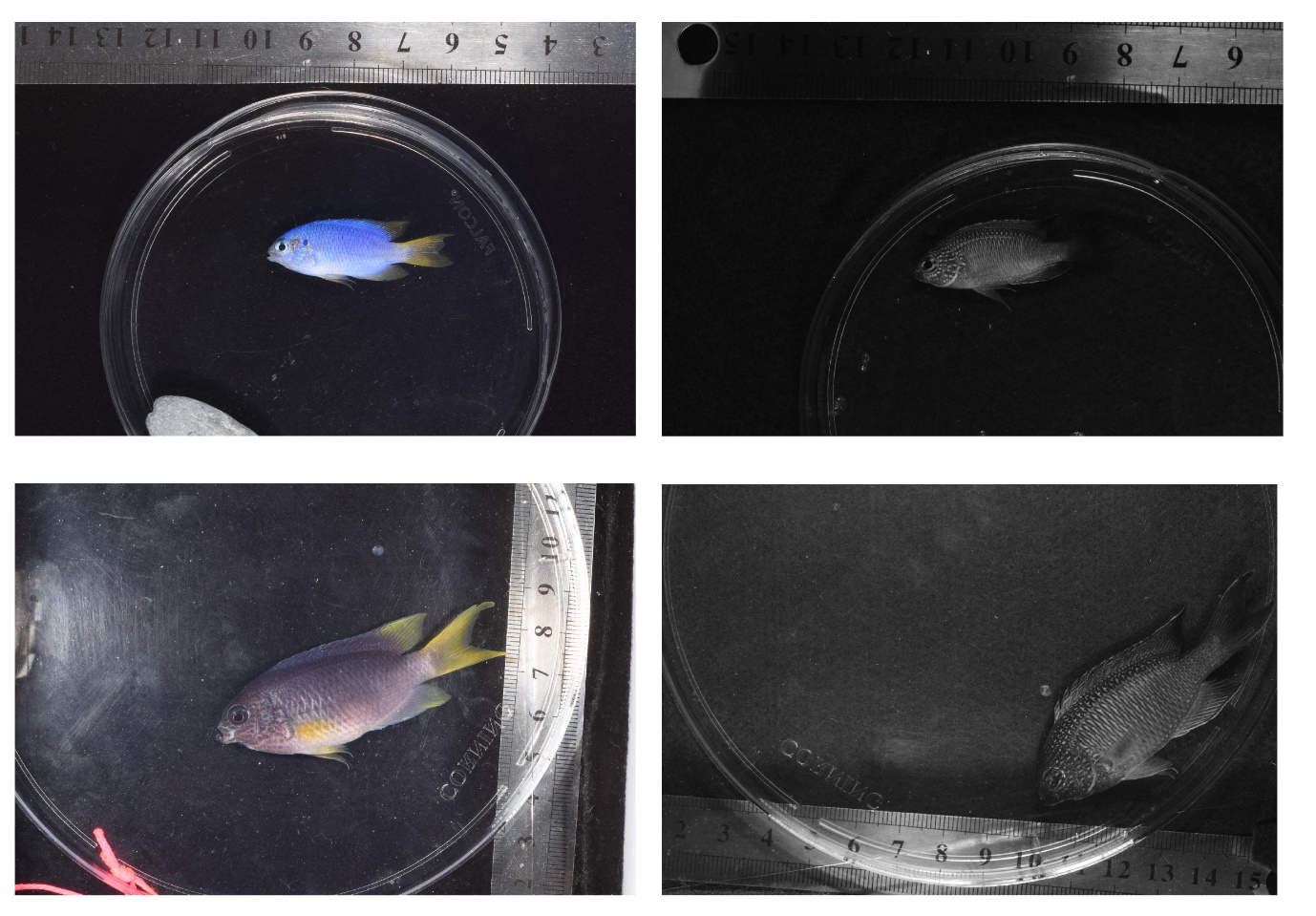

**Figure S16** Pomacentrus pavo pictures taken both with a stock (left) and an ultraviolet camera (right). From the top: juvenile, and adult stages.

Supplementary Methods

PCR for COI amplification

Primers (from Ward et al. 2005):

FishF1 - 5’ TCAACCAACCACAAAGACATTGGCAC 3’

FishF2 - 5’ TCGACTAATCATAAAGATATCGGCAC 3’

FishR1 - 5’ TAGACTTCTGGGTGGCCAAAGAATCA 3’

FishR2 - 5’ ACTTCAGGGTGACCGAAGAATCAGAA 3

Settings:

| Cycle Number | Denature | Anneal | Extend |
| --- | --- | --- | --- |
| 1 | 95 °C, 1 min |  |  |
| 2-36 | 94 °C, 30 s | 56 °C, 45 s | 68 °C, 60 s |
| 37 |  |  | 68 °C, 5 min |
